# A multivariate brain signature for reward

**DOI:** 10.1101/2022.06.16.496388

**Authors:** Sebastian P.H. Speer, Christian Keysers, Ale Smidts, Maarten A.S. Boksem, Tor D. Wager, Valeria Gazzola

## Abstract

The processing of rewards and losses are crucial for learning to adapt to an ever changing environment. Dysregulated reward processes are prevalent in mental health and substance use disorders. While many human brain measures related to reward have been based on activity in individual brain regions, recent studies indicate that many affective and motivational processes are encoded in distributed systems that span multiple regions. Consequently, decoding these processes using individual regions yields small effect sizes and limited reliability, whereas predictive models based on distributed patterns yield much larger effect sizes and excellent reliability. To create such a predictive model for the processes of rewards and losses, from now on termed the Brain Reward Signature (BRS), we trained a LASSO-PCR model to predict the signed magnitude of monetary rewards and losses on the Monetary Incentive Delay task (MID; N = 39) and achieved a high significant decoding performance (92% for decoding rewards versus losses). We subsequently demonstrate the generalizability of our signature on another version of the MID in a different sample (92% decoding accuracy for rewards versus losses; N = 12) and on a gambling task from a large sample (73% decoding accuracy for rewards versus losses, N = 1084) from the Human Connectome Project. Lastly, we also provided preliminary evidence for specificity to rewarding outcomes by illustrating that the signature map generates estimates that significantly differ between rewarding and negative feedback (92% decoding accuracy) but do not differ for conditions that differ in disgust rather than reward in a novel Disgust-Delay Task (N = 39). We thus created a BRS that can be used to make specific, generalizable and reproducible predictions about brain responses to rewards and losses.

## Introduction

The processing of rewards and losses is central to guiding our actions towards positively valenced outcomes and away from negatively valenced ones (Lutz & Widmer, 2014). Numerous functional Magnetic Resonance Imaging (fMRI) studies have investigated the neural correlates of reward processing and several meta-analyses have synthesized the findings of these studies (Bartra et al., 2013; Clithero & Rangel, 2014; Diekhof et al., 2012; Liu et al., 2011). They generally converge on two main insights: First, receiving a reward, or a loss, evokes activity in the nucleus accumbens and surrounding ventral striatum that is hypothesized to represent a positive, or negative, prediction error signal, respectively, defined as the difference between the actual outcome and the one that was expected (Diekhof et al., 2012; Galtress et al., 2012; Haber & Knutson, 2010; O’Doherty et al., 2004). This signal is essential for learning as it increases the likelihood of behavior leading to better than expected outcomes (McClure et al., 2004; Schultz & Dickinson, 2000; Yacubian, 2006) and reduces that of behavior leading to worse than expected outcomes. Second, obtaining abstract goods such as money (but also other categories such as food & nonfood consumables etc. see Chib et al., 2009), recruits the ventro-medial prefrontal cortex (vmPFC) (Kringelbach, 2004; Sescousse et al., 2013), the activity of which is thought to represent the subjective value of a received good (Bartra et al., 2013; Diekhof et al., 2012; Haber & Knutson, 2010; Levy & Glimcher, 2012; Peters & Büchel, 2010) and is also involved in integrating goal information and conceptual information into this value signal (Hare et al., 2008; Plassmann et al., 2007). Using MVPA, McNamee and colleagues (2013) found that spatially distributed patterns in the dorsal part of the vMPFC encodes goal-value information that is independent of stimulus category, whereas the more ventral part of the vmPFC encodes unique category dependent value signals in spatially distinct areas.

Most of these studies have so far used a univariate approach that aims at identifying the locations in the brain recruited while participants process rewards. In some cases, however, the aim is not to map a circuit involved in reward, but to perform reverse inference by asking whether reward processing is involved in a given task X, based on the pattern of brain activity measured at a particular moment in that task (Poldrack, 2006). It has been shown that finding activity in a particular region of the brain is a poor indicator of the recruitment of a particular mental process, because most locations are recruited while engaging a number of mental processes (Poldrack, 2006; Wager et al., 2016). In contrast, a pattern of activity across many voxels, that can include reductions and increases of BOLD signal, has been shown to be associated with a particular mental process with higher sensitivity and specificity, and therefore to provide scientists with a helpful tool to evaluate how strongly a specific mental process is recruited in a given task (Wager et al., 2013; Yarkoni et al., 2011). The ability to decode the degree to which someone is receiving a reward or a loss has yet to be developed. The advantages of such a multivariate brain model are that it leads to larger effect sizes in brain-outcome association compared to more traditional local region-based approaches; makes quantitative predictions about outcomes that can be empirically falsified and can be tested and validated across studies and labs which promotes reproducibility (for a review on brain signatures see Kragel et al., 2018).

Here we therefore aim to develop such a multivariate brain model for reward processing - the brain reward signature (*BRS*) *-* that would use distributed information within and across brain regions to make population-level, between-subject predictions about the strength of engagement of reward processing. These predictions should ideally generalize accurately across contexts, and be able to distinguish reward processing from other categories of related mental processes, such as (emotional) salience (Kragel et al., 2018). So far few signatures for reward-related processes are available (Grosenick et al., 2013) and to our knowledge none of these have been validated on independent samples. A recent large scale challenge to predict Autism Spectrum Disorder diagnoses from fMRI (>146 team & fMRI from > 2000 individuals) highlighted the importance of validating predictive models in independent datasets because model development on a given dataset faces the risk of overfitting. Specifically, techniques such as cross-validation to measure predictive performance are not completely robust to systematic exploration of analytic choices, because the models may overfit on noise that is specific to the data set the models are trained on.

Consequently, our study thus further contributes by validating the *BRS* in three independent samples.

So in this study we use a predictive modeling approach (Kragel et al., 2018) that has been successfully employed to explore the neural representation of various affective processes, including the degree of physical pain (Wager et al., 2013), vicarious pain (Krishnan et al., 2016), social rejection (Woo et al., 2014), unpleasant pictures (Chang et al., 2015), basic emotions (Kragel et al., 2016; Kragel & LaBar, 2015; Lindquist & Barrett, 2012; Saarimäki et al., 2018; Wager et al., 2015), empathy (Ashar et al., 2017), guilt (Yu et al., 2020), and also faces and object categories (Haxby et al., 2001), intentions (Haynes et al., 2007; Soon et al., 2013), semantics (Huth et al., 2012, 2016) and clinical conditions (Arbabshirani et al., 2017; Woo et al., 2017). Our primary goal is to create a *signed relative BRS*. Specifically, the objective is to create a signature that generates more positive values for conditions associated with higher rewards, and more negative values for conditions associated with higher losses.

Additionally, the signature should be specific: it should not generate high pattern responses in datasets in which reward processing should be absent, but other positive or negative emotions were evoked, such as disgust or guilt. Third, it should generalize across studies, samples and contexts where the same neurocognitive processes are engaged (i.e., be generalizable).

Based on our aim to generate a signed relative signature, we trained and tested a LASSO-PCR model (least absolute shrinkage and selection operator-regularized principal components regression; Wager et al., 2011, 2013) to predict the signed magnitude of reward received in the Monetary-Incentive-Delay task (MID, N = 39; see Methods) to establish the *BRS* and test its performance as quantified based on the correlation coefficient between the actual reward value and the pattern response from the neural signature. The pattern response is defined as the dot product between the *BRS* and the parameter estimates from a given condition and task plus the intercept.

The MID was used because it is the most consistently used task to investigate the neural correlates of reward processing in humans (more than 200 MRI studies until now; (Oldham et al., 2018) and has been designed on the basis of findings that reward anticipation engages dopaminergic neurons in the ventral tegmental area (VTA; Knutson et al., 2000). One strength of the MID is that it allows to model a simple decision, which reduces the cognitive confounds that are associated with more complex decision making (Balodis & Potenza, 2015; Knutson & Greer, 2008; Lutz & Widmer, 2014), reliably. Further, the MID robustly engages the striatum, which is crucial in reward processing (Haber & Knutson, 2010). To further probe the performance but also the generalizability, we then applied the *BRS* to a different version of the MID (with 5 instead of three levels of reward; N =12) from different participants using different scanners and scanning protocols (Srirangarajan et al., 2021). Besides that, we also tested the *BRS* in a completely different task with monetary outcomes using a block design instead of an event-related design on a large sample (1084 subjects) to thoroughly evaluate the generalizability of the predictions from our signature map.

Finally, to examine the specificity of the *BRS,* we employed the novel Disgust-Delay Task (DDT, N = 39; Figure 1D), which evokes neural patterns associated with disgust. In this task, we aimed at exploring whether the signature is specific to monetary rewards and losses or rewarding outcomes more generally (i.e. positive versus negative feedback) and whether it is specific to reward or generalizes to emotional salience (i.e. disgusting versus neutral outcomes). The DDT was chosen because it is similar in task structure and solely differs in the neurocognitive processes it is designed to elicit.

**Figure 1.**
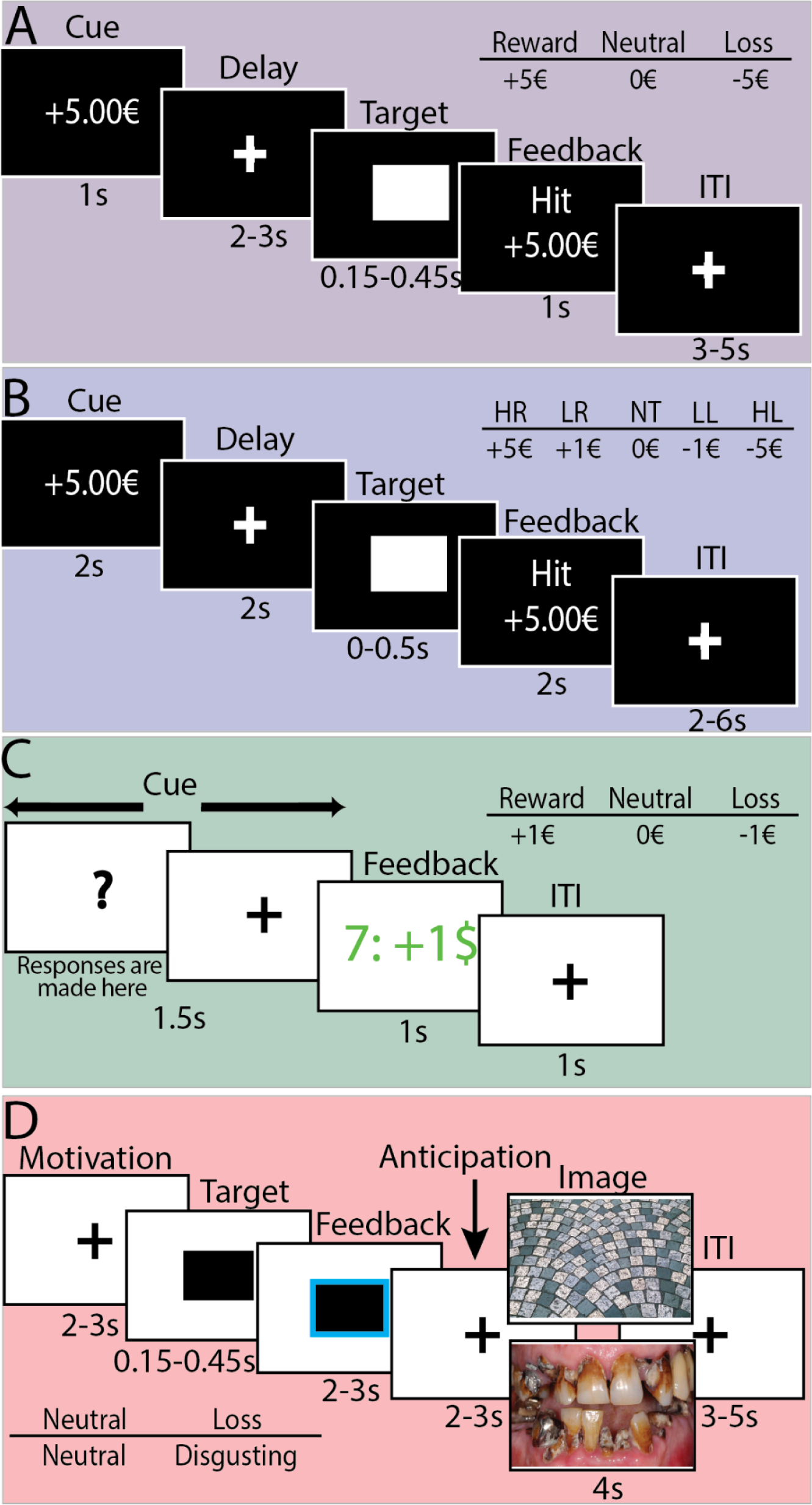
A) Example trial of the MID_train_ task: Each trial started with a cue informing participants about the money that can be obtained or lost. Subsequently, participants were presented with a fixation cross for a variable amount of time (2-3 s) and the target in the form of a white square appeared for a variable amount of time. Afterwards, participants were informed whether they hit or missed and the associated monetary outcome was presented. Lastly, another fixation cross was presented for a variable amount of time (3-5 s). B) Example trial of the MID_val_ task. The differences to the MID_train_ consisted of differences in timing and number of conditions. C) Example trial of the Gambling task from the HCP: Each trial began with the presentation of the mystery card represented by the question mark and as soon as the participants responded, a fixation cross was presented. Next, participants received feedback about the outcome for 1 s. Lastly, another fixation cross was presented for 1 s. D) Example trial of the Disgust-Delay task: Each trial began with the presentation of a fixation cross (2-3 s) followed by a target which was presented for a duration that adapted to the participants’ performance. Next, participants received feedback (2-3 s), viewed another fixation cross (2-3 s) and were then presented with a disgusting or neutral image contingent on their performance. The trials were separated by a fixation cross (3-5 s).

Collectively, datasets from 4 independent studies (N = 1135) were used to train and test the *BRS.* It is important to note that testing specificity is an open ended process, as numerous different conditions unrelated to outcome processing can be tested, but this is a preliminary validation.

As we are interested in investigating the neural underpinnings of reward processing more generally and not the neural correlates of how much exactly someone earns on the MID, our performance assessment focuses on the signature’s relative performance, i.e., whether the signature can predict differences in rewards across conditions. This is because it has been consistently shown across species that value-based choice behavior is context dependent (Bateson et al., 2003; Huber et al., 1982; Shafir et al., 2002; Simonson, 1989). Specifically, it has been found that how a chooser decides between any two options depends on the number or quality of other options in multidimensional attribute space (Huber et al., 1982; Louie et al., 2013). This context-dependence of value based decisions is hypothesized to be implemented on the neural level by means of divisive normalization (Louie et al., 2011, 2013, 2014), where the response of a given neuron is divided by the summed activity of a larger neuronal pool (Carandini & Heeger, 2012). This divisive normalization thus produces context dependence, where the value of an option is explicitly contingent on the value of the other available options, which allows efficient coding of information in changing environments. Therefore, our feature selection procedure was based on correlations between actual and regression-predicted rewards, to capture the relative predictive performance, not the absolute predictive performance. This focus on within-subject differences between conditions also has the advantage to be less sensitive to confounding individual differences such as vascular response properties.

## Methods

For this project, data from four different studies were used. First, to establish the *BRS*, the Monetary-Incentive-Delay task (MID; Figure 1A) with three levels of monetary outcomes (+5 €, 0 €, -5€) was used, which will from now on be referred to as MID_train_. To test whether the *BRS* generalizes to the MID task with five levels of monetary outcomes (+5 €, + 1€, 0 €, -1 €, -5 €) from different participants using different scanners and scanning parameters, openly available data from Srirangarajan and colleagues (2021) was used (Figure 1B). This dataset will from now on be referred to as MID validation task (MID_va)l_. In addition, to investigate whether the *BRS* is able to predict differences in reward in a different task with monetary outcomes using a block design instead of an event related design we utilized the Gambling task (Figure 1C) from the Human Connectome project. Lastly, to assess the construct validity and test whether the signature is specific to monetary reward, and does not generalize to emotional salience, we employed the novel Disgust-Delay Task (DDT; Figure 1D).

### Participants

For the MID_train_ and the DDT task the same 40 participants were used which were collected from a university sample. One participant had a hit rate of zero in both tasks, indicating that the participant never experienced reward. We thus excluded this participant from the analysis. The remaining 39 participants (*M_age_*= 23.62, *SD_age_* = 3.17; 28 females) were right-handed with normal or corrected to normal vision, spoke English fluently, were not on any psychoactive medication influencing cognitive function, and had no record of neurological or psychiatric illness. The study was approved by the Erasmus Research Institute of Management (ERIM; Protocol NR: 2018/02/06-61976ssp) internal review board and was conducted according to the Declaration of Helsinki.

For the MID_val_, nineteen subjects completed the MID task while being scanned with a multi-band acquisition protocol. According to the pre registered exclusion criteria, data from three subjects were excluded due to excessive motion during at least one of the three task runs, while data from four subjects were excluded due to equipment failure (i.e., faulty response registration by a new button box), leaving twelve subjects total for analyses. For the justification of the sample size and details about participants see the paper by Srirangarajan and colleagues (2021) or contact the authors (Srirangarajan and colleagues).

For the HCP gambling task, task-based fMRI recordings were used from 1200 participants (HCP All Family Subjects). Out of these 1200 participants, 1084 had complete fMRI data for both runs of the Gambling task. Additional behavioral and demographic measures on the individual participants can be downloaded from the project website (Van Essen et al., 2012).

### Task and Stimuli

#### MID_train_

The MID_train_ consisted of 108 trials of approximately 9 s each. During each trial, participants saw one of three cues (cue phase, 1 s), were then asked to fixate on a crosshair as they waited a variable interval (delay phase, 2000–3000 ms), and then responded to a white target square that appeared for a variable length of time (target phase, 150–450 ms) with a button press (Figure 1A). Feedback (outcome phase, 1 s), which followed the disappearance of the target, notified participants whether they had won or lost money during that trial. On incentivized trials, participants could win or avoid losing money by pressing the button during target presentation. On neutral trials, no money could be won or lost. Task difficulty, in the form of the length of time the target was presented, was set adaptively throughout the task such that each participant should succeed on 66% of his or her target responses. This was done to make subjects with different performance levels comparable and prevent participants from getting frustrated. Cues signaled potential reward (+ 5.00 €), potential loss ( - 5.00 €), or no monetary outcome (0 €).

Trial types were pseudo-randomly ordered within each session (Knutson et al., 2000). Participants were instructed that at the end of the experiment one trial would randomly be chosen and that the performance on this trial would determine their remuneration. In the MID task we focus on the feedback phase as we are interested in the neural response associated with receiving a monetary outcome.

#### MID_val_

Since the main goal of the study by Srirangarajan and colleagues (2021) was to examine whether acquiring FMRI data with multi-band versus single-band scanning protocols compromises detection of mesolimbic activity during reward processing, the fMRI data was collected in three runs. Importantly, the MID task was identical across all three runs. The MID_val_ was similar to the MID_train_ with some exceptions. First, the MID_val_ included six task trial conditions: a large gain condition (+5.00 $); a medium gain condition (+1.00 $); a no gain condition (+ $0.00); a no loss condition (- $0.00); medium loss condition (– 1.00 $); and a large loss condition (–5.00 $). Each trial condition was repeated 12 times in a pseudorandom order, totalling 72 trials. Furthermore, timing differed slightly. The cue phase was now 0–2 s, the delay phase was 2–4 s, the target phase appeared briefly between 4–4.5 s, the outcome phase lasted 6–8 s, and the Inter-Trial Interval lasted 2, 4, or 6 s. Thus, each trial lasted an average of 12 s (including the ITI). As before, adaptive timing of target duration within condition ensured that subjects succeeded in “hitting” targets on approximately 66% of the trials (Knutson et al., 2005). Thus, each MID task run lasted 864 s in total (approximately 14.4 min), and all three runs were acquired during a single session, but with counterbalanced ordering across subjects.

### Gambling task from the Human Connectome Project (HCP)

This task was adapted from the Gambling task developed by Delgado and Fiez (Delgado et al., 2000). Participants played a card guessing game where they were asked to guess the number on a mystery card (represented by a “?”) in order to win or lose money (Figure 1C). Participants were told that potential card numbers ranged from 1-9 and were asked to indicate whether they expected the mystery card number to be more or less than 5 by pressing one of two buttons on the response box. Feedback was the number on the card generated by the program as a function of whether the trial was a reward, loss or neutral trial, and could result in: 1) a green up arrow with “$1” for reward trials, 2) a red down arrow next to -$0.50 for loss trials; or 3) the number 5 and a gray double headed arrow for neutral trials. The “?” was presented for up to 1500 ms (if the participant responds before 1500 ms, a fixation cross was displayed for the remaining time), followed by feedback for 1000 ms. There was a 1000 ms ITI with a “+” presented on the screen. The task was presented in blocks of 8 trials that are either mostly reward (6 reward trials pseudo randomly interleaved with either 1 neutral and 1 loss trial, 2 neutral trials, or 2 loss trials) or mostly loss (6 loss trials pseudorandomly interleaved with either 1 neutral and 1 reward trial, 2 neutral trials, or 2 reward trials). In each of the two runs, there were 2 mostly reward and 2 mostly loss blocks, interleaved with 4 fixation blocks (15 s each).

This experiment was designed to be analyzed in blocks of mainly reward blocks and mainly loss blocks. As a consequence, here we do not analyze a specific period within each trial, but the average activation across several trials within each block type.

### Disgust-Delay task

A new paradigm termed the Disgust-Delay-Task (DDT) inspired by the monetary incentive delay task(Knutson et al., 2000) was developed (Figure 1D). In this task, participants had to press a button during the presentation of a target stimulus, i.e.,a black rectangle. They were then informed, during the feedback phase, about whether the trial was a success or not. However, instead of winning money, or avoiding losing money, during the outcome phase, participant then either saw a disgusting image or a neutral image depending on their performance. Disgusting images were selected based on a pretest that ensured that these images evoked disgust specifically and no other negatively valenced emotions (see Appendix 1). On each trial of the DDT, participants were first presented with a fixation cross for 2-3s (Figure 1D). Subsequently, the target stimulus was presented for 150-450 ms depending on the participants’ performance. As in the MID tasks above, an adaptive algorithm was implemented which varies the duration to ensure an equal number of successful and unsuccessful trials (50% each). Afterwards, the participants received feedback whether or not they hit the target in time for a period that varied between 2-3 s. This was followed by another fixation cross that varied between 2-3 s. The trial ended with the presentation of either a neutral image or a disgusting image for 4 s depending on whether the participant hit or missed the target. Next, participants had to wait for a period jittered between 3-5 s. Participants completed 72 trials of the DDT. Here, we can thus analyse two periods of interest. During the feedback period, we can investigate the impact of a non-financial reinforcer (i.e., success or failure feedback) on brain activity. During the outcome phase, we can investigate the impact of neural response to the experience of disgust triggered by the disgusting images.

### fMRI acquisition

For MID_train_ and DDT, the fMRI images were collected using a 3T Siemens Verio MRI system. Functional scans were acquired by a T2*-weighted gradient-echo, echo-planar pulse sequence in descending interleaved order (3.0 mm slice thickness, 3.0 × 3.0 mm in-plane resolution, 64 × 64 voxels per slice, flip angle = 75°). TE was 30 ms, and TR was 2,030 ms. A T1-weighted image was acquired for anatomical reference (1.0 × 0.5 × 0.5 mm resolution, 192 sagittal slices, flip angle = 9°, TE = 2.26 ms, TR = 1,900 ms).

For MID_val_, all data were acquired on a 3 Tesla General Electric scanner with a 32-channel head coil at the Stanford Center for Cognitive and Neurobiological Imaging (CNI). Structural (T1-weighted) scans were first acquired for all participants. Functional (T2 ∗ -weighted) images for single-band and multi-band scans were then acquired using the following common parameters: TE = 25 ms, FOV = 23.8 ×23.8 cm; 2 acquisition matrix = 70 ×70, no gap, phase encoding = PA, voxel dimensions = 3.4 ×3.4 ×3.4 mm. Additional parameters that varied between scanning protocols included: (1) multi-band factor = 1, TR = 2000 msec, flip angle = 77°, number of slices = 41; (2) multi-band factor = 4, TR = 500 msec, flip angle = 42°, number of slices = 32; (3) multi-band factor = 8, TR = 500 msec, flip angle = 42°, number of slices = 41. All FMRI data were reconstructed using 1D-GRAPPA (Blaimer et al., 2013). For more information about the scanning protocol please refer to the paper by Srirangarajan and colleagues (2021).

For the HCP project, the data was collected using a customized 3T Siemens Connectome Skyra with a standard 32-channel Siemens receiver head coil and a body transmission coil.

T1-weighted high-resolution structural images were acquired using a 3D MPRAGE sequence with 0.7 mm isotropic resolution (FOV = 224 × 224 mm, matrix = 320 × 320, 256 sagittal slices, TR = 2400 ms, TE = 2.14 ms, TI = 1000 ms, FA = 8◦) and used to register functional MRI data to a standard brain space. Functional MRI data were collected using gradient-echo echo-planar imaging (EPI) with 2.0 mm isotropic resolution (FOV = 208 × 180 mm, matrix = 104 × 90, 72 slices, TR = 720 ms, TE = 33.1 ms, FA = 52◦, multiband factor = 8, 253 frames, ∼3 m and 12 s/run).

### Preprocessing

For the MID_train_, MID_val_ and the DDT, the fMRI data were preprocessed using fMRIPrep version 1.0.8, a Nipype based tool (Gorgolewski et al., 2011). We chose fMRIPrep because it addresses the challenge of robust and reproducible preprocessing as it automatically adapts a workflow based on best-in-class algorithms to virtually any dataset, enabling high-quality preprocessing without the need of manual intervention (Esteban et al., 2019). Each T1w volume was corrected for intensity nonuniformity and skullstripped. Spatial normalization to the International Consortium for Brain Mapping 152 Nonlinear Asymmetrical template version 2009c (Esteban et al., 2016) was performed through nonlinear registration, using brain-extracted versions of both T1w volume and template. Brain tissue segmentation of cerebrospinal fluid (CSF), white matter (WM), and gray matter was performed on the brain-extracted T1w. Field map distortion correction was performed by coregistering the functional image to the same-subject T1w image with intensity inverted (Caballero-Gaudes & Reynolds, 2017) constrained with an average field map template (Tustison et al., 2010). This was followed by coregistration to the corresponding T1w using boundary-based registration (Smith et al., 2002) with 9 degrees of freedom. Motion correcting transformations, field distortion correcting warp, blood oxygen level-dependent images-to-T1w transformation, and T1w to template Montreal Imaging Institute (MNI) warp were concatenated and applied in a single step using Lanczos interpolation. Physiological noise regressors were extracted using CompCor (Cox & Hyde, 1997). Principal components were estimated for the two CompCor variants: temporal (tCompCor) and anatomical (aCompCor). Six tCompCor components were then calculated including only the top 5% variable voxels within that subcortical mask. For aCompCor, six components were calculated within the intersection of the subcortical mask and the union of CSF and WM masks calculated in T1w space, after their projection to the native space of each functional run. Frame-wise displacement (Treiber et al., 2016) was calculated for each functional run using the implementation of Nipype. For more details of the pipeline, see https://fmriprep.org/en/latest/workflows.html. After the preprocessing the voxel size of the images is 3*3*3.5 mm.

For the HCP data, Preprocessing of the images included motion correction, distortion correction, co-registration and normalized to MNI space as described in the HCP 1200 Subjects Release (Glasser et al., 2013).

### Statistical analyses

*MID_train_* & *MID_val._* To model all possible outcomes of the MID tasks for every participant, we estimated a general linear model (GLM) using regressors for onsets of the outcome phase for successful high reward trials (HR-won: received + 5.00 €), unsuccessful high reward trials (HR-lost: did not receive +5.00 €), successful low reward trials (LR-won: received + 1.00€; for MID_val_ only), unsuccessful low reward trials (LR-lost: did not receive + 1.00€; for MID_val_ only), successful neutral trials (NT-won: 0 €; for the MID_val_ the neutral gain, i.e.. +0 €, and neutral loss trials, i.e. -0 € were combined), unsuccessful neutral trial (NT-lost: 0€), successful low loss trials (LL-won: did not lose 1.00 €; for MID_val_ only), unsuccessful low loss trials (LL-lost: did lose 1.00 €; for MID_val_ only), successful high loss trials (HL-won: did not lose 5.00€) and unsuccessful high loss trials (HL-lost: lost 5.00€). The duration of the epoch for the outcome phase was 1 s, and the beginning of the outcome phase was used as onset time. Average background, WM and CSF signal, framewise displacement, six head motion regressors, and six aCompCor (which are component based noise correction regressors) regressors, all obtained from fMRIprep, were entered as regressors of no interest. First, a smoothing kernel of 5 mm full width at half maximum was applied. For consistency, the same smoothing procedure was applied to all other datasets as well. Subsequently, all regressors of interest (but not regressors of no interest) were convolved with the canonical hemodynamic response function. Linear contrasts were computed between HR-won and HR-lost trials, LR-won and LR-lost trial, NT-won and NT-lost trials, LL-lost and LL-won trial, HL-lost and HL-won trials. These contrasts were chosen to isolate the effect of receiving or losing money by means of comparing each regressor with the regressor of opposite outcome within the same condition. As a consequence, only neural activation related to receiving or losing money should remain as all other aspects of the contrasted trials are the same. The resulting subject level t-maps were then converted to z-maps. Here, we use the z-maps as the primary input to our multivariate pattern analysis because z-maps represent effect-sizes in units of variance, that should be more comparable across experiments and designs than the simple difference between the parameter estimates, which are in arbitrary units, or the t-maps that depend on the sample size in terms of acquired volumes. As the purpose of the study by Srirangarajan and colleagues (2021) was to test whether acquiring FMRI data with multi-band versus single-band scanning protocols compromises detection of mesolimbic activity during reward processing, the fMRI data was collected in three runs. For this study we were however not interested in the effects of scanning protocols. As a consequence, we averaged over the z-maps for each subject across the three runs to increase the signal to noise ratio.

*DDT.* To model the experience of disgust and the experience of viewing neutral images we estimated a GLM using regressors for onsets of the picture presentation phase of the DDT for the presentation of disgusting images and neutral images. The duration of the epoch for the picture presentation phase was 4 s, and the beginning of the picture presentation phase was used as onset time (see Figure 1D).

In addition, to explore whether the *BRS* predicts *monetary* outcomes specifically or generalizes to rewarding versus loss outcomes more generally, we modeled the feedback phase of the DDT. As the structure of the MID and the DDT are very similar the only difference here is that instead of monetary outcome the feedback is purely motivational. The duration of the epoch for the feedback phase was 2 s since this was the minimum of time it lasted on every trial. We defined the feedback phase by counting back two seconds from the onset of the Anticipation phase (see Figure 1D). Lastly, to have a neutral period to compare the neural patterns associated with disgusting and neutral images to, we modeled the neural activation of viewing the fixation cross at the beginning of each trial (Motivation Delay). This period was chosen because it was most distant in time from the picture presentation phase. The duration of the epoch for the motivation delay was 2 s since this was the minimum of time it lasted on every trial (see Figure 1D). As above, average background, WM and CSF signal, framewise displacement, six head motion regressors, and six aCompCor regressors, all obtained from fMRIprep, were entered as regressors of no interest. First, a smoothing kernel of 5 mm full width at half maximum was applied. Next, all regressors of interest (but not the nuisance regressors) were convolved with the canonical hemodynamic response function. Linear contrasts were computed between the presentation of a disgusting images and the fixation period and the presentation of a neutral image and the fixation period. As before, the subject level t-maps were converted to z-maps to render them more comparable across experiments.

*HCP.* Since the HCP gambling task was administered in a block design and the ITIs between trials were short we employed a GLM using regressors for onsets of the reward blocks, loss blocks and fixation blocks. The duration of the reward and loss blocks were 28s each whereas the fixation period was 15s. Twelve motion regressors (x translation in mm, y translation in mm, z translation in mm, x rotation in degrees, y rotation in degrees, z rotation in degrees, derivative of x translation, derivative of y translation, derivative of z translation, derivative of x rotation, derivative of y rotation, derivative of z rotation), the absolute root mean square (RMS) motion and the relative RMS motion, obtained from the HCP preprocessing pipeline, were added as regressors of no interest. Different nuisance regressors were applied here as the data was obtained in preprocessed format from the HCP website and only the 14 regressors mentioned in the previous sentence were available. As before, as a first step, a smoothing kernel of 5 mm full width at half maximum (FWHM) was applied. Afterwards, all regressors of interest (but not the regressors of no interest) were convolved with the canonical hemodynamic response function.

Linear contrasts were computed between the reward block and the fixation block, the loss block and the fixation block and the fixation block and the baseline. Again, the resulting subject level t-maps were subsequently converted to z-maps.

### Multivariate pattern analyses

#### Creation of the BRS

We used the normalized and smoothed (5mm FWHM) z-maps to develop population-level reward-predictive patterns, as previous studies suggested that smoothing could improve inter-subject functional alignment while retaining sensitivity to mesoscopic activity patterns that are consistent across subjects (Etzel et al., 2011; Op de Beeck, 2010; Shmuel et al., 2010). A LASSO-PCR model (least absolute shrinkage and selection operator-regularized principal components regression; Wager et al., 2011, 2013) was then trained on the whole-brain maps from the subject level z-maps derived from the analyses described above. Specifically, the LASSO-PCR model was trained on the z- maps (HR-won > HR-lost, NT-won > NT-lost; HL-lost > HL-won) from the MID_train_ to predict the 3 different levels of monetary outcome (+ 5.00 €, 0.00 € & -5.00€). For feature selection, we identified voxels that correlated more strongly with reward rather than salience. As explained in the introduction, this was done to maximize relative prediction performance rather than absolute prediction, because reward processing has been found to be context dependent (Bateson et al., 2003; Huber et al., 1982; Louie et al., 2013; Shafir et al., 2002; Simonson, 1989) and there are no absolute values assigned to individual options. Specifically, given the three parameter estimate images for each participant (High Reward: HR-won > HR-lost, Neutral: NT-won > NT-lost, High Loss: HL-lost > HL-won), we can consider two codings: one for outcome (1, 0, -1) and one for salience (1, 0, 1). We can then compute the Spearman correlation between the parameter estimates *V_j_* at each voxels *j* and the outcome and salience coding separatly for each subject within the cross validation loop. As we know that the spacing is uncertain, because rewards might not be equidistant from zero as losses (Kahneman, 2011), we use the Spearman instead of the Pearson correlation. We then selected voxels such that 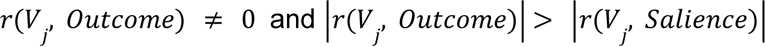. At the group level, to do this, we first performed a two-sided Wilcoxon signed-rank test on the correlation between voxel values and outcome coding *r*(*V*_*j*’_, *Outcome*) and then a one-sided Wilcoxon signed-rank test on the difference between absolute values of the correlation between voxel values and outcome and voxel values and salience 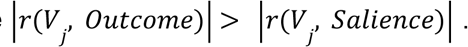 We then selected all voxels for which 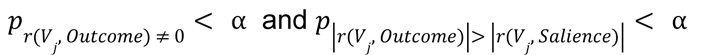 where α was chosen permissively at α=0.5 to allow for a reasonable amount of voxels to enter the LASSO-PCR model. More conservative thresholds were also applied to test the robustness of the findings (see Appendix 2). To reiterate the feature selection procedure, we correlated for each subject the parameter estimates for each of the three conditions (High Reward, Neutral & High Loss) with the two codings (outcome and salience) at each voxel, to select the voxels that correlate more strongly with the outcome coding than with the salience coding, while making sure that the voxels respond to the outcome coding. This was done on each iteration of the cross-validation on the training set to only allow voxels to enter the LASSO-PCR model that respond stronger to outcome than to salience.

The feature selection and model fitting were implemented using a 5-fold cross-validation procedure during which all participants were randomly assigned to 5 different subsamples while ensuring that all images from an individual subject remained within a subsample and does not spread across subsamples. We always used 4 subsamples for training and one for testing. As a result, out-of-sample prediction is always done on new individuals, which prevents dependence across images from the same participants invalidating predictive accuracy. To evaluate the predictive accuracy of the model the Spearman correlation between the predicted monetary outcome levels and the actual outcomes for the left-out subsample were computed at each fold and then the correlations were averaged across folds. In accordance with the mass-univariate analyses and to identify which brain regions made reliable contributions to the model (Wager et al., 2013; Zhou et al., 2020), the pattern maps were thresholded at p < 0.001 (two-tailed; uncorrected) using bootstrap procedures with 5000 samples. The result was a spatial pattern of regression weights across the whole brain that significantly contributed to the prediction of monetary out-of-sample outcomes in the MID_train_. To test for robustness, we also applied a more conservative threshold at FDR p < 0.05 (two-tailed) and a procedure in which we first selected only voxels that were non-zero in at least 90% of the bootstrap iteration and then applied FDR correction at q< 0.05 (see Appendix 2). We also computed the Bayes-Factor for the correlation between predicted and actual monetary outcome values to also be able to test for the evidence for the absence of an effect (Keysers et al., 2020). To calculate the Bayes-Factor for the correlation Jeffreys exact Bayes Factor was used (Ly et al., 2016) as implemented in the Pingouin python package (Vallat, 2018). In addition, we evaluated whether the *BRS ’s* predictions within a given condition (High Reward, Neutral, High Loss) are significantly different from zero, by means of a one sample t-test against zero. Since not all of the predictions across conditions and experiments were normally distributed we used the Wilcoxon signed-rank test and the associated Bayes factors were computed as proposed by van Doorn, Marsman and Wagenmakers (2020), with a Cauchy prior with the scale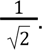. To compare the the *BRS ’s* predictions between conditions Wilcoxon rank-sum tests were employed and to compute the Bayes Factors we again used the procedure proposed by van Doorn and colleagues (2020).

We also conducted within-person forced-choice discrimination, where two activation maps from the same participant were compared, and the image with the higher overall signature response (i.e., the stronger expression of the signature pattern) was classified as associated with higher reward. We conducted these forced-choice tests for all combinations of conditions (i.e., HR vs. NT, NT vs HP & HR vs HP). The advantage of the forced-choice test is that it is ‘threshold free’ in the sense that an absolute decision threshold across individuals is not required; zero is used as the threshold for the difference between the two paired alternatives (Wager et al., 2013).

Thus, individual differences in the shape and amplitude of the blood oxygen level dependent (BOLD) fMRI response do not add noise in this kind of test. To test for significance permutation tests were used where the order of conditions was permuted (N = 10000) and the accuracy was computed again. The empirical classification accuracy was then compared to the null distribution of accuracies based on permuted values to obtain p-values.

#### Validation on the MID_val_

To test how well the *BRS* generalizes to new data involving monetary outcomes the *MID_val_* was used. Specifically, we tested whether the *BRS* generalizes to a MID task with five levels of monetary outcomes (+5 €, + 1€, 0 €, -1 €, -5 €) from different participants using different scanners and scanning parameters. To this end, we obtained pattern expression values by computing the dot product of the cross-validated weightmap (averaged across folds) of the reward pattern (created on the MID_train_) and the z-maps and adding the intercept (averaged across folds) for each subject and condition from the MID_val_. For the MID_val_ the High Reward (HR-won > HR-lost), Low Reward (LR-won > LR-lost), Neutral (NT-won > NT-lost), Low Loss (LL-lost > LL-won) and High Loss (HL-lost > HL-won) contrasts were used. The resulting pattern expression represents scalar response values, which constitute the predicted monetary outcome for the given condition. The pattern expression values were then tested for differences between experimental conditions. We calculated the Spearman correlation between the pattern expression values and the actual monetary outcome values for each of the conditions (+5 €, + 1€, 0 €, -1 €, -5 €), with higher correlations representing higher predictive accuracy, in the sense of variance of rewards explained by the pattern expression values. Specifically, the predicted monetary outcome values obtained from the dot multiplication (5 conditions * 12 subjects = 60 predicted monetary outcome values) were correlated with the 60 actual monetary outcomes in a single correlation. To estimate significance of the predictive performance, a permutation test (N = 5000) was performed where the true monetary outcome values were shuffled and the procedure was repeated. To assess the robustness of the estimation of significance we also repeated the permutation tests with the root mean squared error as an predictive performance evaluation metric (N = 5000). To test whether the predictions made by the *BRS* in the different conditions were different from zero and whether predictions between conditions were significantly different from each other, the same procedure as detailed above was used. As before, we conducted within-person forced-choice discrimination, to further assess the predictive accuracy of the *BRS.* As above, permutation testing was used to evaluate statistical significance of classification accuracies.

### Validation on the HCP Gambling task

To investigate whether our *BRS* generalizes to a completely different task involving monetary outcomes the HCP Gambling task was used. Specifically, we tested whether the *BRS* generalizes to the Gambling with three different levels of monetary outcomes (+ 1€, 0 €, -0.5 €) that were not symmetrically distributed around zero. As before, we obtained pattern expression values by computing the dot product of the cross-validated weightmap (averaged over folds) of the reward pattern (created on the MID_train_) and the z-maps and adding the intercept (averaged over folds) for each subject and condition from the HCP Gambling task and then tested the predictive performance using the Spearman correlation between actual monetary outcomes and predicted monetary outcome values (3 conditions * 1084 subjects = 3252 predicted monetary outcome values). As above, permutation tests were used to estimate significance.To test whether the predictions made by the *BRS* in the different conditions were different from zero and whether predictions between conditions were significantly different from each other, the same procedure as detailed above was used. Again, we conducted within-person forced-choice discrimination, to further assess the predicitive accuracy of the *BRS.* As above, permutation testing was used to evaluate statistical significance of classification accuracies. To evaluate the test-retest reliability of the HCP Gambling task, we also computed the pattern response to the first and second run separately and then calculated the pearson, spearman and intraclass correlation between the pattern responses for the two runs. We chose to assess test-retest reliability for the HCP specifically because it was the only sample large enough to get meaningful estimates of test-retest reliability.

### Testing the specificity on the DDT task

For specificity, the signature expression should not significantly differ from zero when applied to z-maps from tasks involving other types of emotionally salient outcomes. To assess the specificity of our *BRS* we employed the DDT task. We explored whether the *BRS* also predicts disgusting (coded as -1) versus neutral outcomes (coded as 0). In addition, we also tested whether the *BRS* would be able to predict positive or negative feedback in the disgust delay task. This was done to explore whether the *BRS* predicts *monetary* outcomes specifically or generalizes to rewarding versus loss outcomes more generally. As before, we obtained pattern expression values by computing the dot product of the cross-validated weight map (averaged over folds) of the reward pattern (created on the MID_train_) and the z-maps and adding the intercept (averaged over folds) for each subject and condition from the DDT task and then tested the predictive performance using the Spearman correlation between actual emotional outcomes (neutral vs disgusting images) and predicted emotional outcomes (2 conditions * 39 subjects = 78 predicted emotional outcome values. As above, permutation tests were used to estimate significance.To test whether the predictions made by the *BRS* in the different conditions were different from zero and whether predictions between conditions were significantly different from each other, the same procedure as detailed above was used. As above, we conducted within-person forced-choice discrimination, to further assess the prediciive accuracy of the *BRS.* As above, permutation testing was used to evaluate statistical significance of classification accuracies.

## Results

### Within-task prediction

To create a generalizable *BRS* we first trained and tested our LASSOPCR model on the MID_train_ using 5-fold cross validation and a threshold of p < 0.5 (threshold was applied within the cross-validation loop) for the feature selection procedure. The analysis revealed that outcomes in the left-out cross-validation folds in the MID_train_ could be significantly predicted by the *BRS* (*RMSE* = 2.89, *p_perm_* < 0.001, *r* = 0.72, *p_perm_*< 0.001, BF_10_ > 1000). The feature selection procedure selected 39% of voxels across the whole brain (Figure 2A). Using the bootstrap procedure, we observed that particularly voxels in the dorsal striatum and the ventromedial prefrontal cortex (vmPFC) significantly contributed to the predictive success of our model (at p < 0.001; Figure 2B and Table 2; for other thresholds see Appendix 2). Figure 3A shows the signature values obtained when multiplying the z-maps of the individual participants with the thresholded (p_bootstrap_ < 0.001; see methods) *BRS*. For the forced choice analysis we observed significant classification accuracies for all tests. However, classification accuracy was substantially higher between rewarding and loss conditions and neutral and loss trials than between reward and neutral conditions (see Table 3).

**Figure 2.**
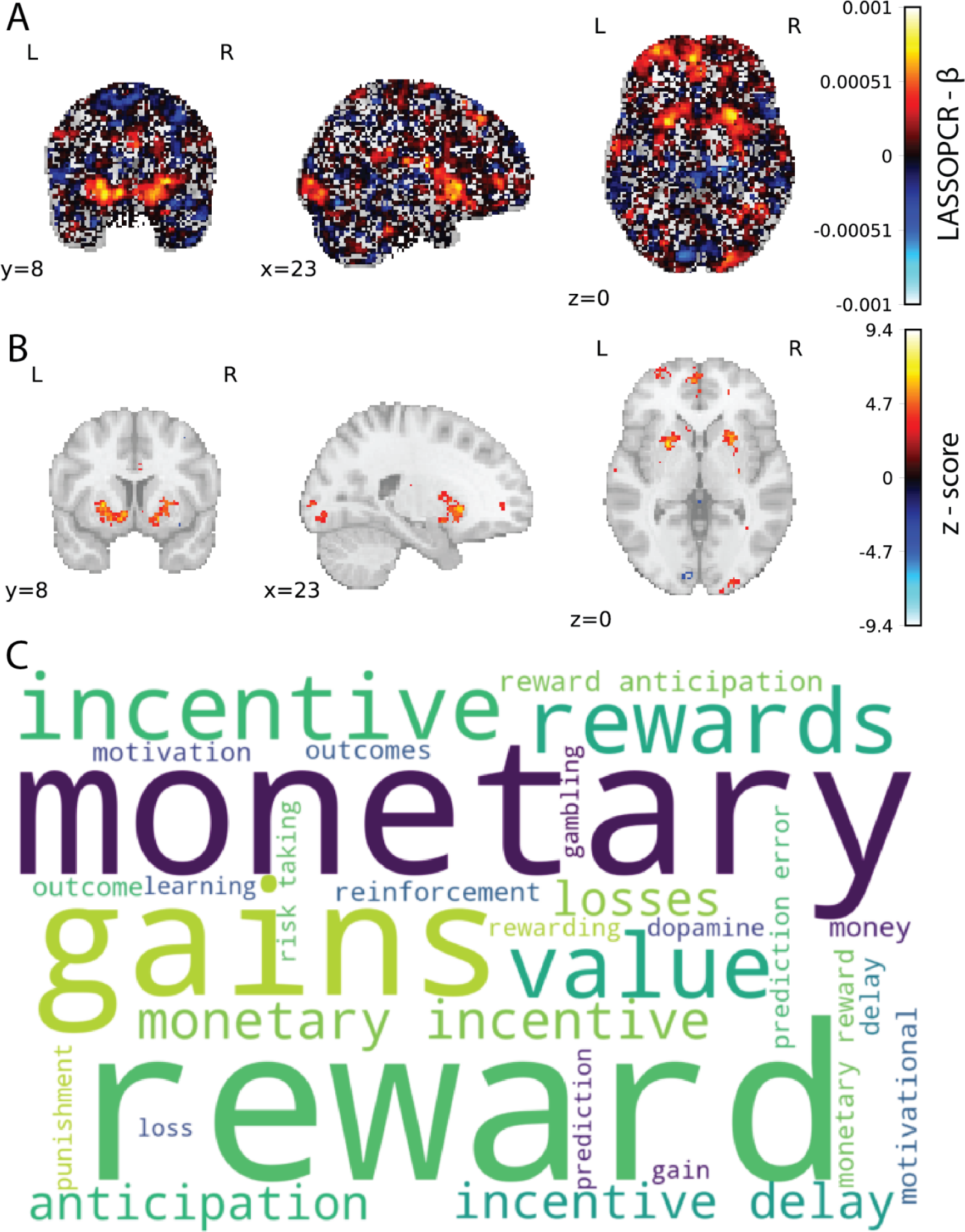
A: Mean weights for the out-of sample prediction on the MID_train_. B: Voxels significantly contributing to the out-of-sample prediction identified using the bootstrap procedure (p<0.001). **C**: Word cloud showing the top 50 relevant terms (excluding anatomical terms) for the meta-analytic decoding of the *BRS* map. The size of the font was scaled by correlation strength (r_min_ = 0.11, r_max_ = 0.22).

**Figure 3:**
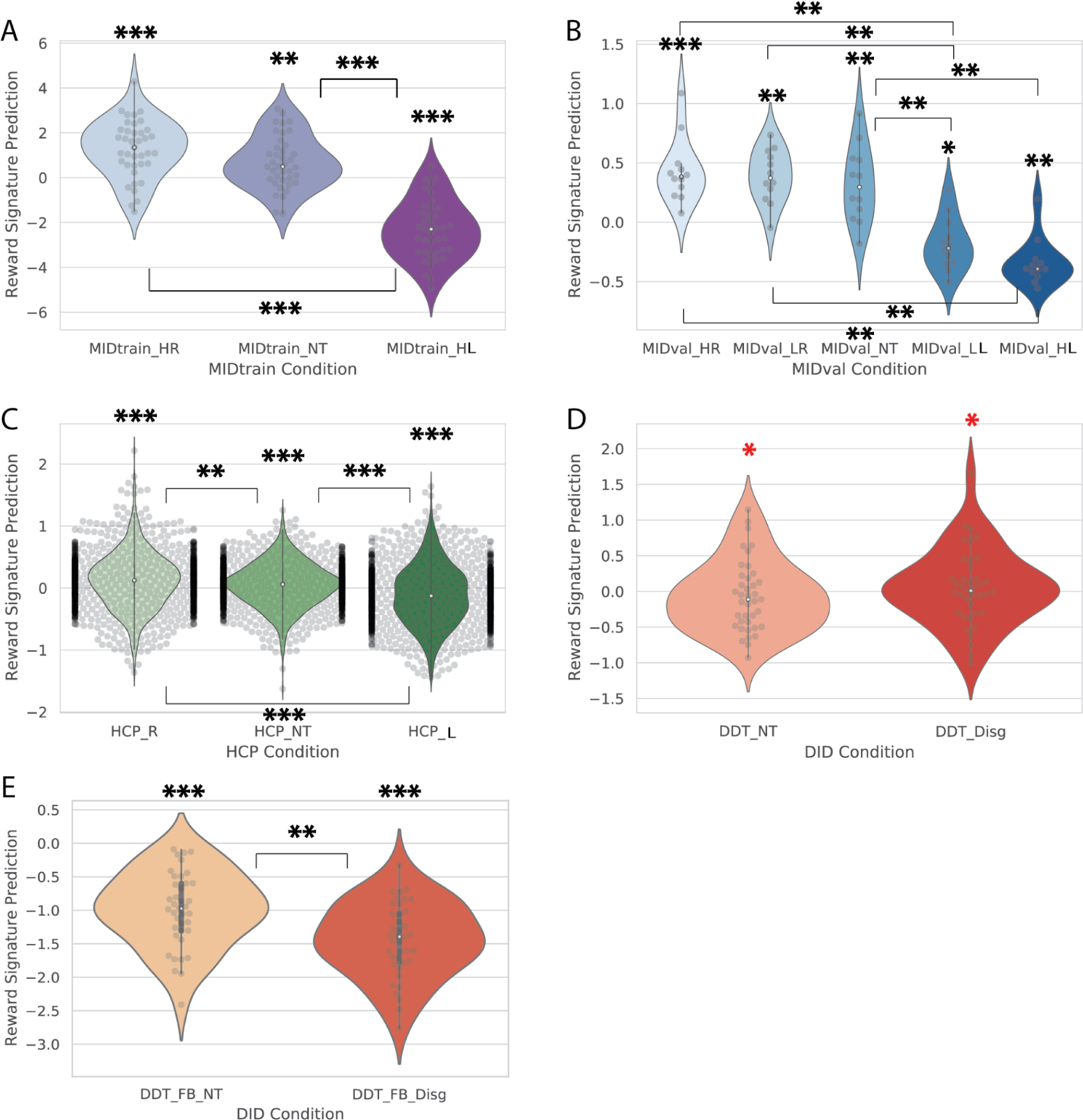
A: Violinplot for the predicted monetary outcomes across conditions in the MID_train._ B: Violinplot for the predicted monetary outcomes across conditions in the MID_val._. C: Violinplot for the predicted monetary outcomes across conditions in the HCP gambling task. D: Violinplot for the predicted disgusting versus neutral outcomes in the DDT for the outcome phase. E: Violinplot for the predicted positive versus negative feedback in the DDT for the feedback phase.. In the violin plot the grey circles represent individual observations arranged so that they do not overlap. The white point represents the median, the top of the box represents the 3rd quartile and lower end of the box represents the first quartile. The top of the upper whisker represents the maximum value and the bottom of the lower whisker represents the minimum value. Because the HCP data contained 1084 subjects only 30% of actual data points could be plotted. The black dots at the edges are due to several points overlapping with each other. DID = Disgust Incentive Delay Task; MID = Monetary Incentive Delay Task; HCP = Human Connectome Project Gambling Task; HR = High Reward; LR = Low Reward; NT = Neutral; LP = Low Loss; HR = High Loss; **\*:**BF_10_ > 3; **\*\*:**BF_10_ > 10; **\*\*\*:**BF_10_ > 100; **\*:**BF_10_ < 0.33. Note that we use BF10 values rather than p-values for the stars in the figure to more balancedly provide evidence for the null or alternative hypothesis. The stars above the violin represents the BF obtained from one-sample t-tests against zero, whereas the stars above the bars between violins represents the BFs obtained from Wilcoxon rank-sum tests comparing predictions between conditions.

**Table 1.** Coding of outcome and salience for feature selection.

| Parameter estimate image | Outcome | Salience |
| --- | --- | --- |
| High Reward (HR) | 1 | 1 |
| Neutral (N) | 0 | 0 |
| High Loss (HL) | -1 | 1 |

**Table 2.**
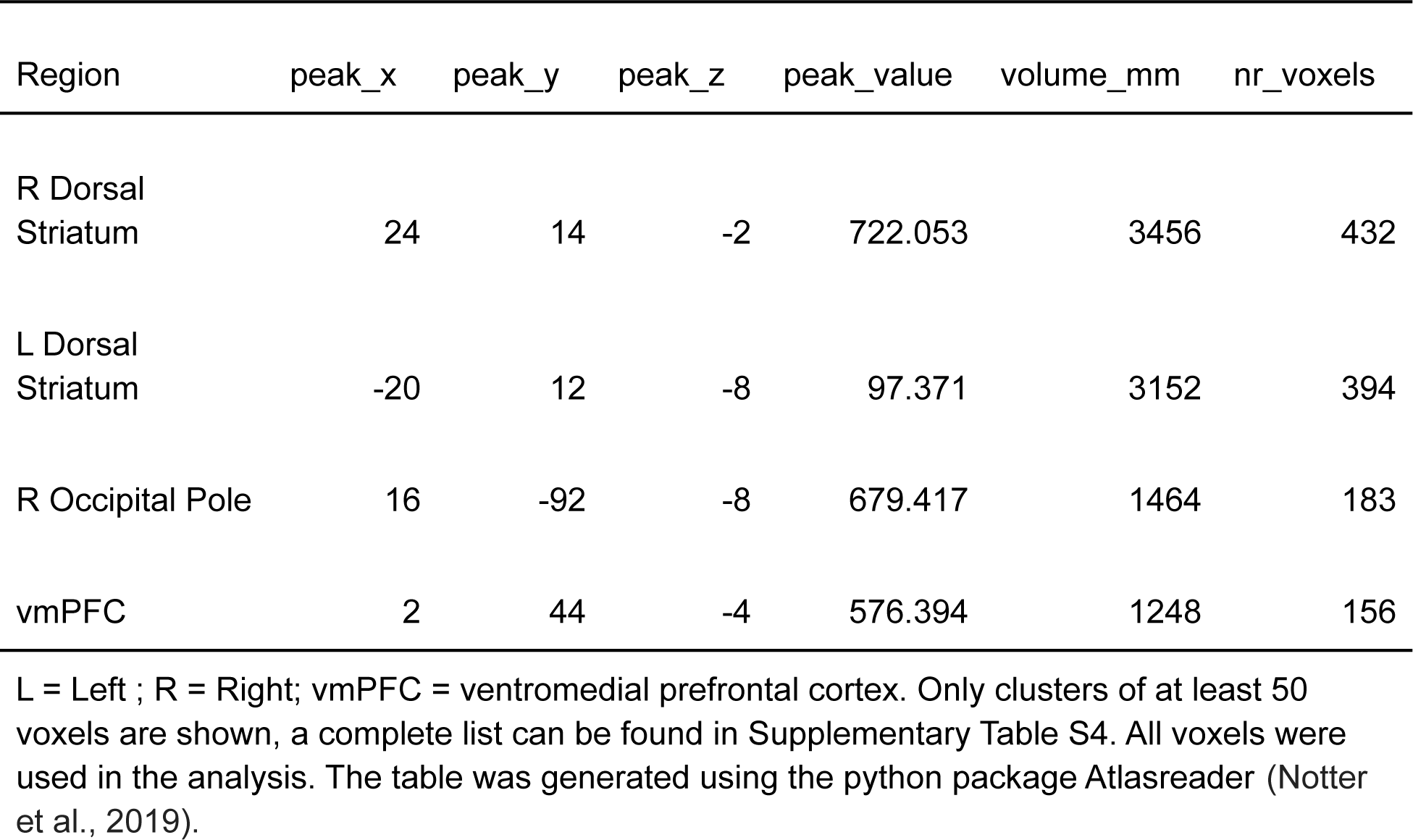
clusters for the significant voxels identified by the bootstrap procedure.

**Table 3.** Forced Choice Accuracies (%) for the MID_train_

|  | HR | N | HL |
| --- | --- | --- | --- |
| HR | - |  |  |
| N | 67*** | - |  |
| HL | 92*** | 95*** | - |

HR = High Reward, N = Neutral; HL = High Loss;
\*= $p_{\text{perm}} < 0.05$ ; \*\*= $p_{\text{perm}} < 0.01$ ; \*\*\*= $p_{\text{perm}} < 0.001$

### Meta-analytic decoding of the BRS map

To functionally characterize the *BRS*, the Neurosynth (Yarkoni et al., 2011) decoder function was used to assess its similarity to the reverse inference meta-analysis maps generated for the entire set of terms included in the Neurosynth dataset.

Here the unthresholded z-map obtained through the bootstrap procedure was used, since the neurosynth decoder works best on unthresholded whole brain maps. The most relevant features were ‘reward’ and ‘monetary’ for the top 50 terms (excluding anatomical terms) ranked by the correlation strengths between the *BRS* map and the meta-analytic maps (see word cloud, size of the font scaled by correlation strength, Figure 2C).

### Testing the generalizability on the MID_val_

To test the generalizability of the *BRS* map we tested the prediction performance on the MID_val_. This allowed us to evaluate how well the *BRS* is able to predict relative reward magnitude based on activation patterns in new participants from a different scanner and with a different number of levels of monetary outcomes. Using the significant voxels from the *BRS* map in Figure 2B we observed a significant prediction of the relative monetary outcomes on the MID_val_ (*RMSE* = 2.97, *r* = 0.75, *p_perm_*< 0.001, BF_10_ > 1000; Figure 3B). To test the robustness of this finding the prediction was also repeated using all voxels, and the FDR-corrected map (q<0.05; see Appendix 2), and a map derived from first selecting the most consistent voxels and correcting using FDR (see Methods). The robustness checks revealed very similar significant predictions on the MID_val_ (see Appendix 2). For the forced choice analysis we observed significant classification accuracies for the tests comparing the rewarding to the loss condition and the neutral to the loss condition. No significant classifcation accuracies were observed when contrasting rewarding and neutral trials. In addition, no significant classification accuracy was observed when comparing high and low loss trials (see Table 4).

**Table 4.**
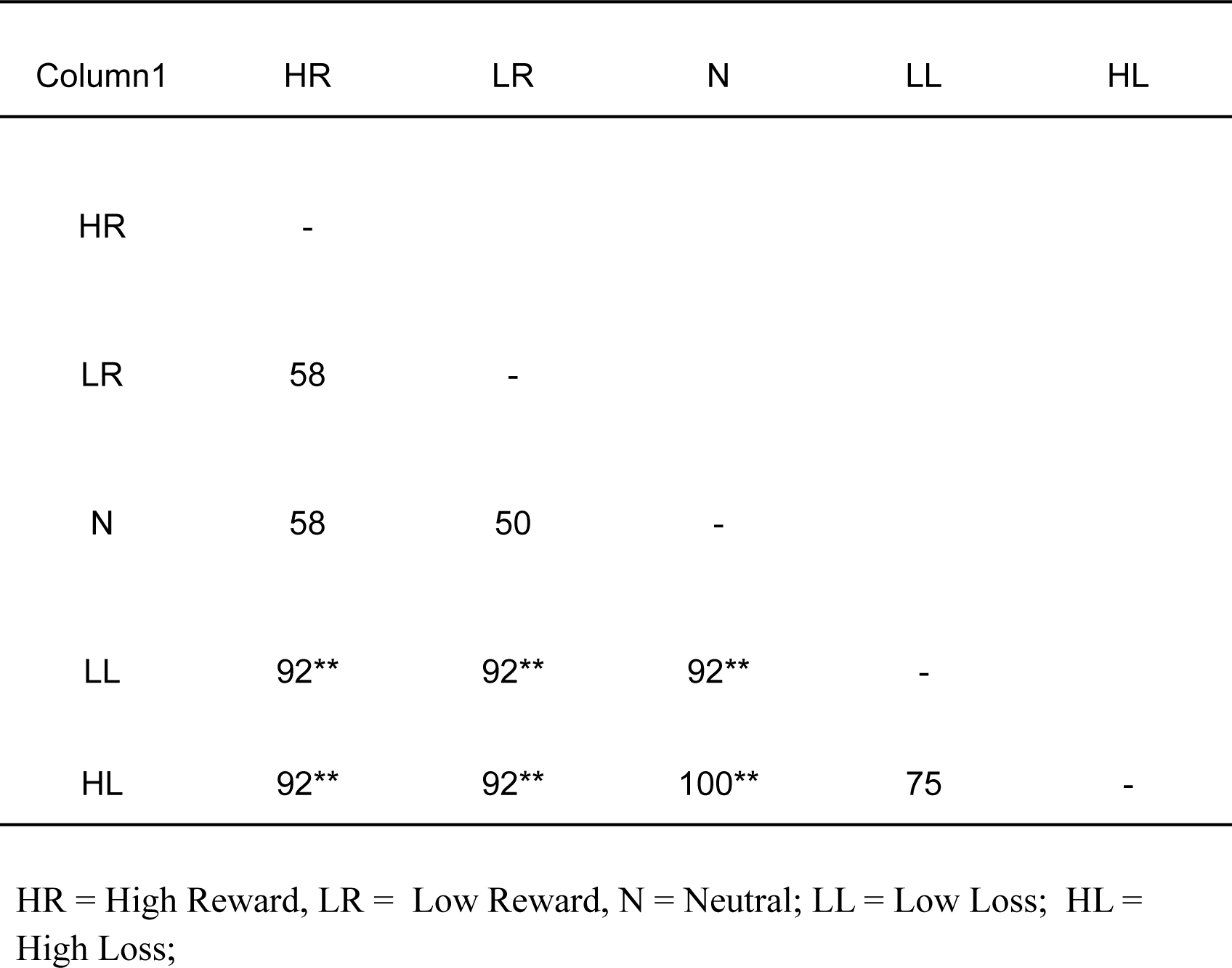
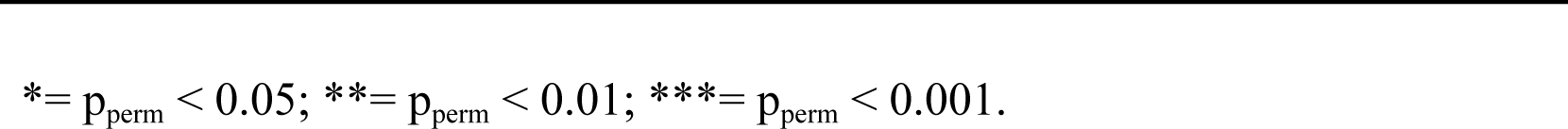
Forced Choice Accuracies (%) for the MID_val_

| Column1 | HR | LR | N | LL | HL |
| --- | --- | --- | --- | --- | --- |
| HR | - |  |  |  |  |
| LR | 58 | - |  |  |  |
| N | 58 | 50 | - |  |  |
| LL | 92** | 92** | 92** | - |  |
| HL | 92** | 92** | 100** | 75 | - |

HR = High Reward, LR = Low Reward, N = Neutral; LL = Low Loss; HL = High Loss;
\*= $p_{\text{perm}} < 0.05$ ; \*\*= $p_{\text{perm}} < 0.01$ ; \*\*\*= $p_{\text{perm}} < 0.001$ .

### Testing the generalizability on the HCP gambling task

To further test the generalizability of the *BRS* map we assessed the prediction performance on the HCP gambling task. This enabled us to test how well the *BRS* is able to predict on a much larger set of participants, from a different scanner, on a different task using a different experimental design (block vs event-related) and with a different asymmetric levels of monetary outcomes. Using the significant voxels from the *BRS* map shown in Figure 2B, we observed a significant prediction of the monetary outcomes on the HCP gambling task (*RMSE* = 0.7,*p_perm_*< 0.001, *r* = 0.21, *p_perm_*< 0.001, BF_10_ > 1000; Figure 3C). To test the robustness of this finding the prediction was also repeated using all voxels, and FDR-corrected map (p<0.05) and a map derived from first selecting the most consistent voxels and the correcting using FDR (see Methods). The robustness checks revealed very similar significant prediction on the HCP gambling task (see Appendix 2). For the forced choice analysis we observed significant classification accuracies for all tests. However, as for the MID tasks, the classification accuracy was substantially higher between rewarding and loss trials and neutral and loss trials than between reward and neutral trials (see Table 5). The analysis of the test-retest reliability revealed that there is a significant correlation between the patterns responses of the first and the second run of the HCP (*r_pearson_* = 0.24, *p_perm_*< 0.001; *r_spearman_* = 0.23, *p_perm_*< 0.001; *r_ICC_* = 0.24, *p_perm_*< 0.001; BF_10_ > 1000).

**Table 5.** Forced Choice Accuracies (%) for the HCP

|  | HR | N | HL |
| --- | --- | --- | --- |
| HR | - |  |  |
| N | 53** | - |  |
| HL | 73*** | 63*** | - |

HR = High Reward, N = Neutral; HL = High Loss; $p_{\text{perm}} < 0.05$ ; \*\*= $p_{\text{perm}} < 0.01$ ; \*\*\*= $p_{\text{perm}} < 0.001$ .
\*=

### Testing the specificity on the DD

In order to evaluate the specificity of the *BRS* map we assessed the prediction performance on the outcome phase of the DDT, in which participants see disgusting or neutral images. This enabled us to investigate whether the *BRS* map predicts differences in emotional salience more generally or whether it more specifically captures differences in reward. Using the significant voxels from the *BRS* map (see Figure 2 middle) we did not observe a significant prediction of the differences in outcomes in the DDT, and most importantly, found Bayesian evidence for the absence of such differentiation (*RMSE* = 0.9, *p_perm_*=0.84, *r* = -0.13, *p_perm_*= 0.28, BF_10_ =0.23; Figure 3D). To test the robustness of this finding the prediction was also repeated using all voxels, and FDR-corrected map (p<0.05), and a map derived from first selecting the most consistent voxels and the correcting using FDR (see Methods). The robustness checks did not reveal any significant prediction on the DDT either (see Appendix 2). For the forced-choice analysis we found that the neutral trials could not be significantly distinguished from disgusting trials in the outcome phase (33%, p = 0.98).

To further assess the specificity of the *BRS* we also tested the feedback phase of the DDT (see Figure 1D), in which participants are informed whether they successfully performed the task or not. Using the significant voxels from the *BRS* map shown in Figure 2B, we found a significant prediction of feedback in the DDT (*RMSE* = 0.92, *p_perm_*< 0.001, *r* = 0.38, *p_perm_*< 0.001, BF_10_ > 1000; Figure 3E). To test the robustness of this finding the prediction was also repeated using all voxels, and FDR-corrected map (p<0.05) and a map derived from first selecting the most consistent voxels and the correcting using FDR (see Methods). The robustness checks revealed very similar significant prediction on the feedback phase of the DDT (see Appendix 2). The forced-choice analysis revealed that the successful trials could be significantly discriminated from unsuccesful trials in the feedback phase (92%, p < 0.001).

## Discussion

In the current study, we developed a multivariate brain model, the *BRS,* that allows us to decode the *relative* degree of reward across conditions. In particular, using the correlation between actual and decoded reward in the MID and HPC gambling task, we show the ability of the *BRS* to explain a significant proportion of the variance in the reward magnitude involved. This *BRS* is not only able to predict variance in the monetary outcome in unseen subjects from the same sample but also generalizes to different samples using a different version of the same task and also to entirely different tasks. Further, this signature was found to not only predict monetary outcomes, but also rewarding outcomes in the form of positive versus negative feedback more generally. Crucially, this *BRS* was found to be specific to rewarding outcomes and did not generalize to emotionally salient (disgusting) images. We thus provide a *BRS* that can be used to make generalizable inferences about the presence of rewarding vs loss outcomes, which does not generalize to a negative but disgusting outcome.

To create the *BRS* that is sensitive to the neurocognitive underpinnings of reward processing, we trained a LASSOPCR model on the MID, which is the most consistently used task to evoke the neural mechanisms associated with processing monetary outcomes (Oldham et al., 2018). To ensure that the *BRS* predicts reward specifically and not salience in general, we only selected voxel for prediction that correlated more strongly with outcome (i.e., voxels that differentiate between reward, neutral and loss outcomes), than with salience (i.e., voxels that differentiate only between neutral and consequential,reward or loss, outcomes). We found that clusters of voxels in the bilateral dorsal striatum, the vmPFC and the right occipital pole significantly decoded monetary outcomes in novel participants from the same sample. We subsequently tested whether the observed clusters indeed reflect reward processing areas by means of using the Neurosynth (Yarkoni et al., 2011) decoder. This decoder compared our *BRS* to the entire set of terms included in the Neurosynth database and found that the highest ranked associations were *reward* and *monetary*, providing converging evidence that the *BRS* predicts rewarding outcomes.

The finding that activation patterns in the dorsal striatum are predictive of rewarding outcomes aligns well with previous fMRI studies, that found that the striatum encodes the prediction error signal (Diekhof et al., 2012; Galtress et al., 2012; Haber & Knutson, 2010; O’Doherty et al., 2004). The striatum has been consistently linked to both the anticipation and evaluation of rewarding outcomes (for review see Oldham et al., 2018). In addition, abnormal activity in the striatum and connectivity between the striatum and the limbic system have been linked to impaired reward processing in obesity and bipolar disorder (Caseras et al., 2013; Nummenmaa et al., 2012; Yip et al., 2015). Similarly, the observation that a cluster of voxels in the vmPFC is predictive of rewrarding outcomes is in accordance with previous fMRI research on economic decisions and reward processing, as it has been associated consistently with the receipt of reward or loss and the computation of subjective value (Bartra et al., 2013; Diekhof et al., 2012; Haber & Knutson, 2010; Kringelbach, 2004; Levy & Glimcher, 2012; Peters & Büchel, 2010; Sescousse et al., 2013). It is relevant to note that while we found reward to be positively associated in our *BRS*, this does not preclude the existence of circuits and ensembles that encode loss and aversive processes and conversely exhibit decreased activation in response to reward.

As a next step, we tested the generalizability of the *BRS* on two different samples. Firstly, we tested the relative predictive accuracy of the *BRS* on a different version of the MID, with five levels of monetary outcomes instead of three, from a different sample and found that we could again decode monetary outcomes significantly with high accuracy, as assessed using the correlation between decoded and actual reward magnitude. Secondly, we assessed the predictive performance of the *BRS* on a large sample (N = 1084) with a different task, namely a gambling task from the Human Connectome Project. Again, we found that the *BRS* was able to significantly predict monetary outcomes. Together, these results highlight the generalizability of the predictions of the *BRS.* The observation that predictive accuracy dropped in comparison to the other two samples can be explained by the fact that this task differed from the MID task in two ways: In contrast to the MID, the gambling included rewards that were not symmetrically distributed around zero. In addition, the gambling task was developed for analysis using a block design (averaging over several trials of the same condition) whereas the MID used an event related design (modeling specific phases within a trial individually).

While our feature selection procedure, which removed voxels that primarily responded to salience, and training the *BRS* on a well-established reward processing task provided a good fundament for ensuring the specificity of predictions, we also wanted to empirically test this specificity. To this end, we also evaluated the predictions of the *BRS* on two phases of the DDT, a novel task designed to evoke disgust as a negative outcome. First, we tested predictions during the feedback phase which provided a success/failure feedback to the participants, and could therefore be triggering neurocognitive processes associated with reward/loss that is non-monetary in nature. Here we found a significant predictive performance of the *BRS*, suggesting that the *BRS* is able to decode reward and loss processing more generally and is not limited to monetary outcomes alone. Second, we tested the outcome phase to test whether predictions are specific to reward or generalize to other emotionally salient outcomes such as disgust. The analysis provided evidence in favor of the absence of an effect. Stated differently, the *BRS* generated predictions that did not differ between participants viewing a disgusting image or a neutral image. This finding suggests that the *BRS* predicts relative rewarding outcomes (financial or otherwise) with some specificity that does not generalize to the other emotionally salient outcome we tested.

Since previous research suggested that reward may be encoded specifically in the striatum (Haber & Knutson, 2010; Knutson et al., 2001, 2005), we also tested whether a broader circuit (i.e. including the vMPFC) is needed to decode reward (see Appendix 3). To this end we applied a more theory driven approach where we used a meta-analytic map created based on the term *monetary reward* and on the term *outcome.* These maps only included voxels in the striatum (and not in the vMPFC). Similar to the data-driven feature selection approach reported in the main text, the *BRS* significantly predicted monetary outcomes in the MID_val_ and the HCP gambling task, and DDT feedback phase, but did not significantly predict outcomes in the DDT outcome phase (see Appendix 3). Performance on the HCP and DDT feedback phase was higher for this theory driven approach as compared to the data driven *BRS*. In contrast, performance on the MID_val_ was slightly lower for the theory driven approach, which was expected as the data-driven *BRS* involved feature selection trained on a version of the MID task similar in nature and consequently was more likely to perform higher on a similar task. In contrast, the theory-driven approach was more task independent and more likely to perform similarly well across tasks. This aligns well with the notion that the striatum may encode reward and losses quite generally.

To test the robustness of our findings, we also repeated all the reported analyses in the main text, using different thresholds for the feature selection procedure and for the correction for multiple comparison (see Appendix 2). These robustness checks validated the findings from the main text. For all feature selection and multiple comparison correction thresholds, the predictions within the MID_train_, MID_test_ , gambling task and feedback phase of the DDT remained significant. Only on the DDT outcome phase (testing for specificity for reward processing), when using all voxels instead of selecting only voxels that were significant in predicting monetary outcomes on the MID_train_, there was not enough evidence to support the hypothesis that the *BRS* was unable to differentiate between disgusting and neutral images. This may be due to voxels contributing to the prediction that are not specific to predicting monetary outcome but also encode emotional salience in general. Since we used a lenient threshold for the feature selection algorithm some voxels coding for salience may have been included in the model and thus lowered the evidence in favor of the absence of an effect. This finding suggests that users should use the signature only including significant voxels when applying the *BRS* to other sets.

In future studies, this *BRS* could be employed to differentiate and compare the contribution of various emotions and cognitive processes to complex (social) decisions. For instance, in the case of moral decisions, it is frequently the case that selfish motives related to monetary benefits are pitted against the concern for others, for example in terms of avoiding harm to a confederate. In such a context, the *BRS* could be applied in combination with neural signatures for vicarious pain (Caspar et al., 2020; Krishnan et al., 2016; Zhou et al., 2020) and for guilt (Yu et al., 2020) to disentangle the contribution of these processes to the eventual decision.

One limitation of the *BRS* so far is that we only tested the specificity of its prediction on a single experimental paradigm with negative emotional salience. To further characterize the specificity of the *BRS* it would be beneficial to test its prediction on experimental paradigm involving positively valenced emotional stimuli, such as for instance snack foods or funny, entertaining or erotic pictures and videos.

A second, critical limitation of our signature to consider when interpreting applications of our *BRS*, is that while its expression values correlate with the reward outcome obtained by participants in the MID and gambling task, the *BRS* failed to identify neutral outcomes as such. Specifically, in the MID tasks (MID_train_ and MID_val_), while we correctly find the gain conditions to generate values significantly above zero, and the loss conditions to generate negative values, the neutral conditions generate values that are also slightly positive (Figure 3a,b). In addition,the BRS failed to discriminate high and low reward conditions in the MID_val_ and also did not differentiate the two rewarding conditions from the neutral condition accurately. Conceptually, this may be explained by the observation that people generally seem to be risk averse (i.e., losses loom larger than gains; Kahneman, 2011). Thus, a loss of the same (monetary) value as a reward will be experienced as more severe and may thus be encoded as more distant from zero (the neutral condition) than a reward of an equal amount. This would explain why our algorithm was not always able to significantly discriminate rewarding from neutral trials, but always achieved significant discrimination between neutral and loss trials. Methodologically, this observation may be explained by the fact that when training a linear model on a dependent variable with only three levels the model will be mostly influenced by its extreme points, whereas the middle point will be less influential in determining parameter estimates. Further, our feature selection algorithm was designed to maximize *relative* prediction performance rather than absolute prediction, because value based computations and associated outcome processing have been found to be context dependent (Bateson et al., 2003; Huber et al., 1982; Louie et al., 2013; Shafir et al., 2002; Simonson, 1989) and that decisions do not reflect absolute valuations assigned to individual alternatives. This however means that our signature should not be applied to the z-values of a single condition to determine if any reward processing was triggered, but rather on multiple conditions to test whether they differ in reward processing.

A last limitation pertains to the fact that several constructs related to reward processing have been associated with the striatum and vMPFC contained in our *BRS*, such as the outcome value, anticipated outcome, goal value and prediction error (Diekhof et al., 2012; Galtress et al., 2012; Haber & Knutson, 2010; Knutson et al., 2005; O’Doherty et al., 2004; Rutledge et al., 2010) and we can’t precisely disentangle which of these processes are captured specifically by our signature. Future studies may aim at more clearly separating the neural signatures of each of these constructs.

In summary, we created a *BRS* that robustly predicts monetary outcomes that generalizes across tasks and several large samples. This *BRS* is specific to rewarding outcomes and does not appear to generalize to at least one other salient emotional outcomes. The benefit of this signature over the univariate approach is that it integrates distributed information from regions across the whole brain into a single optimized prediction which can then be tested across conditions on new and independent individuals and samples. As a consequence, this approach circumvents the need for multiple comparisons and provides unbiased estimates of effect size (Reddan et al., 2017) when assessing the involvement of reward processes in different experimental conditions. This renders the signature approach more sensitive, generalizable and reproducible than traditional univariate approaches (Kragel et al., 2018).

### Data availability

The unthresholded *BRS* can be found on neurovault: https://neurovault.org/images/775976/. The thresholded map and scripts used in the manuscript can be found on Github: https://github.com/SebastianSpeer/Reward_Signature.

Data and scripts used in the task will be made available on OSF.

## Acknowledgements

The authors would like to thank the editor and anonymous reviewers for their insightful comments. We also gratefully acknowledge financial support from the Erasmus Research Institute of Management (ERIM).

## Appendix

### Appendix 1: Pretest for disgusting Images for the DDT

In order to test whether the pictures used to evoke disgust were in fact perceived as disgusting a pretest of the images was conducted. 50 disgusting images (depicting rotten food, insects etc.) were downloaded from the internet. To ensure that pictures were selected that elicited particularly disgust and no other negative emotion we also added images that evoke other negative emotions for comparison. This was achieved by selecting the 50 images that scored highest on negative valence and arousal from the OASIS picture set (Kurdi et al., 2017). A sample of 101 workers from Amazon’s Mechanical Turk then rated all these 100 images on how strongly 8 different emotions, namely Amusement, Awe, Contentment, Excitement, Anger, Disgust, Fear and Sadness (Zhao et al., 2014), were evoked on a rating scale from 0 (*not at all*) to 10 (*extremely strong*) . Based on these rating we then calculated the Euclidean distance from the ideal image (10 on disgust and 0 and all other dimensions) and then standardized the scores (divided each score by the maximum distance) and reverse coded it (1-Euclidean distance) so that the higher the number the closer the images are to the ideal score. The 45 images that ranked highest were selected for the experiment. For the neutral images, 45 images from the OASIS pictures set were selected based on how close they were to the middle point for valence (3.5 on a 7 point Likert-scale from *very negative* to *very positive*) and to the lowest point for arousal (1 on a 7 point Likert-scale from *very low* to *very high*).

### Appendix 2: Robustness checks for the validation tasks

To test the robustness of our LASSOPCR model in prediction on the validation tasks, we repeated the analyses with different thresholds for the feature selection procedure . Thus, we selected all voxels for which 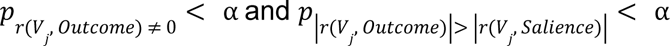 where α was also chosen at α = 0. 3 and α = 0. 4. In addition, we also tested the robustness of our findings by testing different thresholding techniques for the bootstrapped weights. Specifically, we used all voxels, the voxels that survived an FDR threshold at p < 0.05, and voxels that were not zero in at least 90% of bootstrap iterations and survived the FDR threshold at p < 0.05.

Collectively the results closely mirror the findings reported in the main text. We find significant predictions in the MID_val_ and the HCP gambling task, whereas no significant predictions are found for the DDT (see Table S1).

**Table S1.**
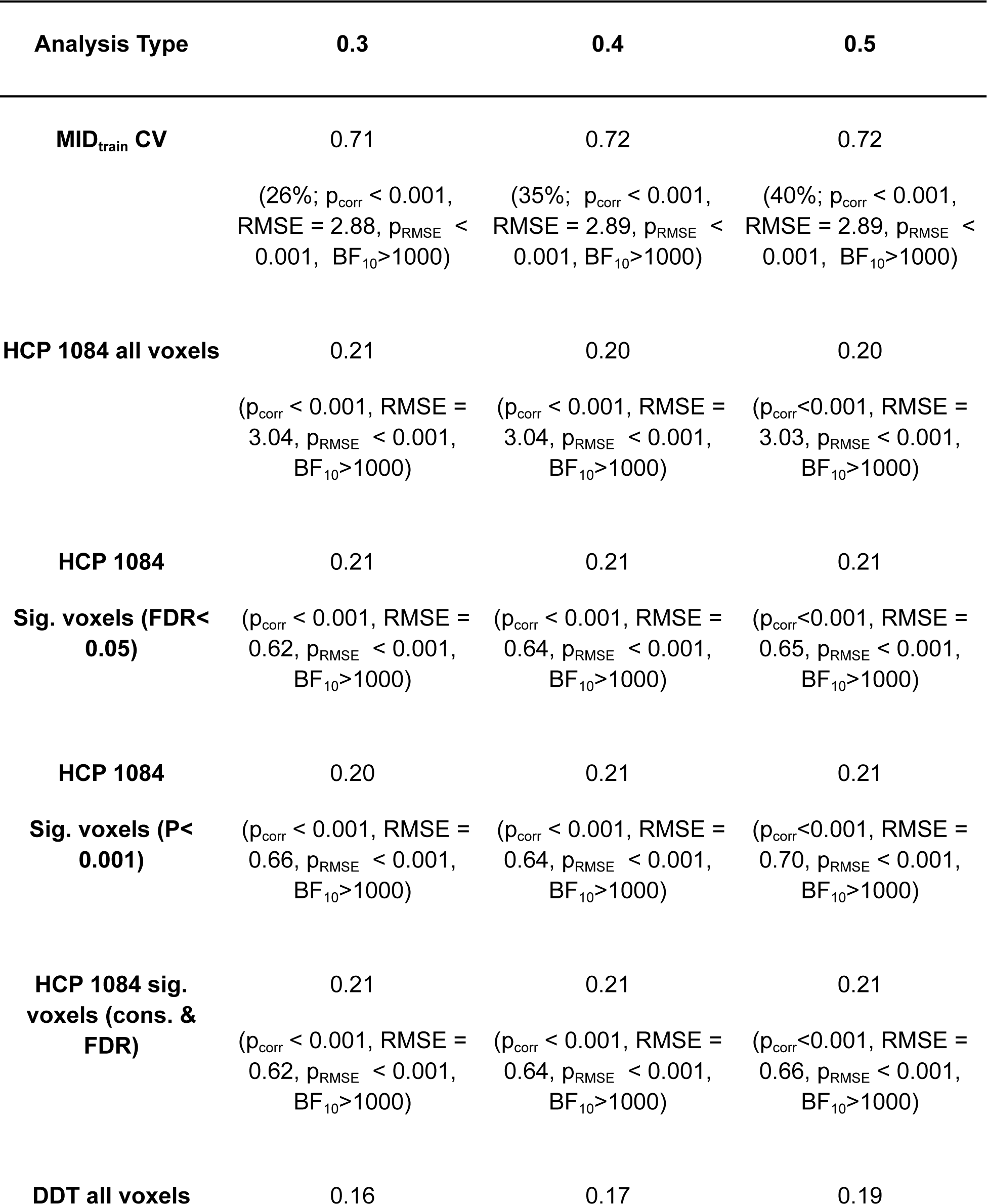
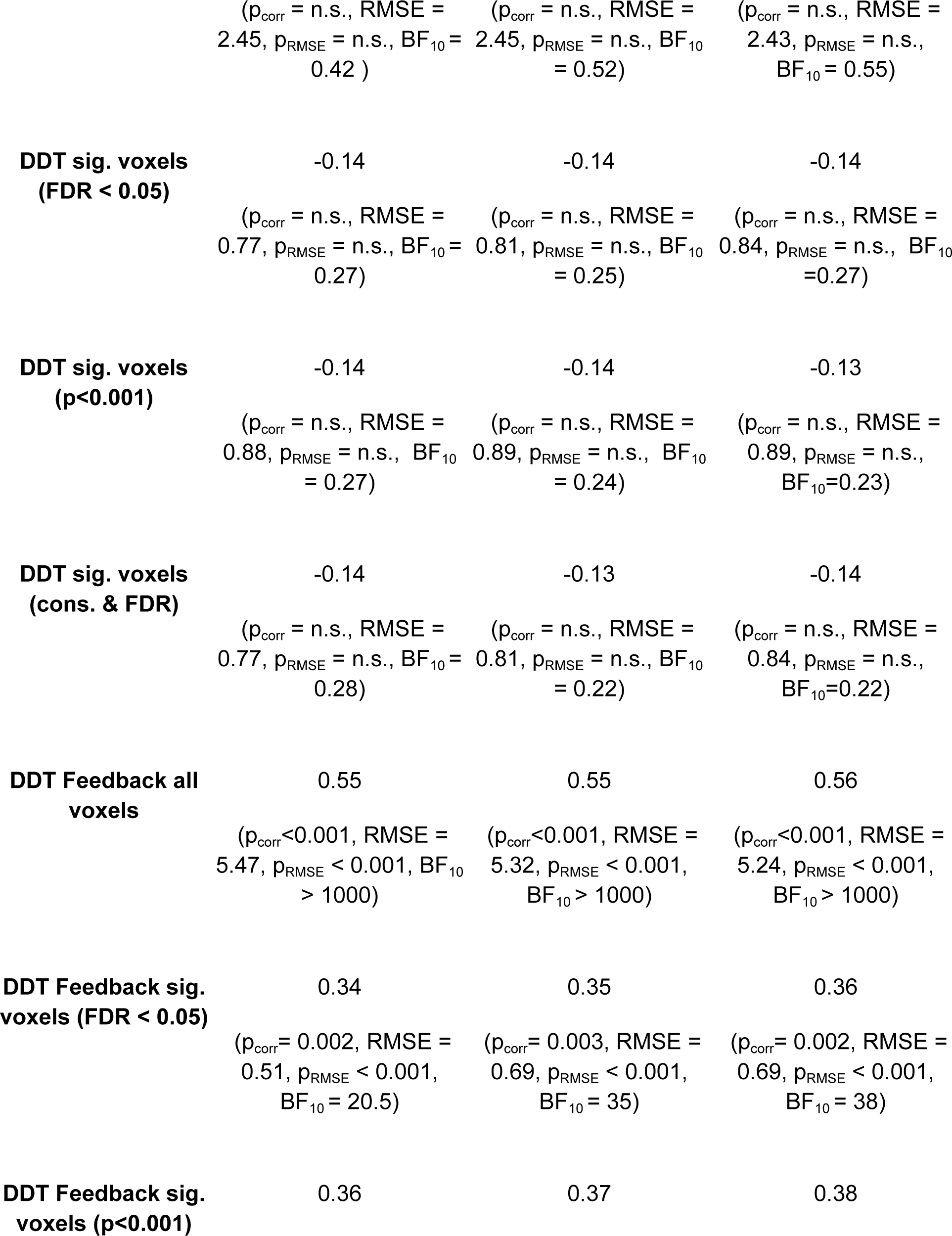
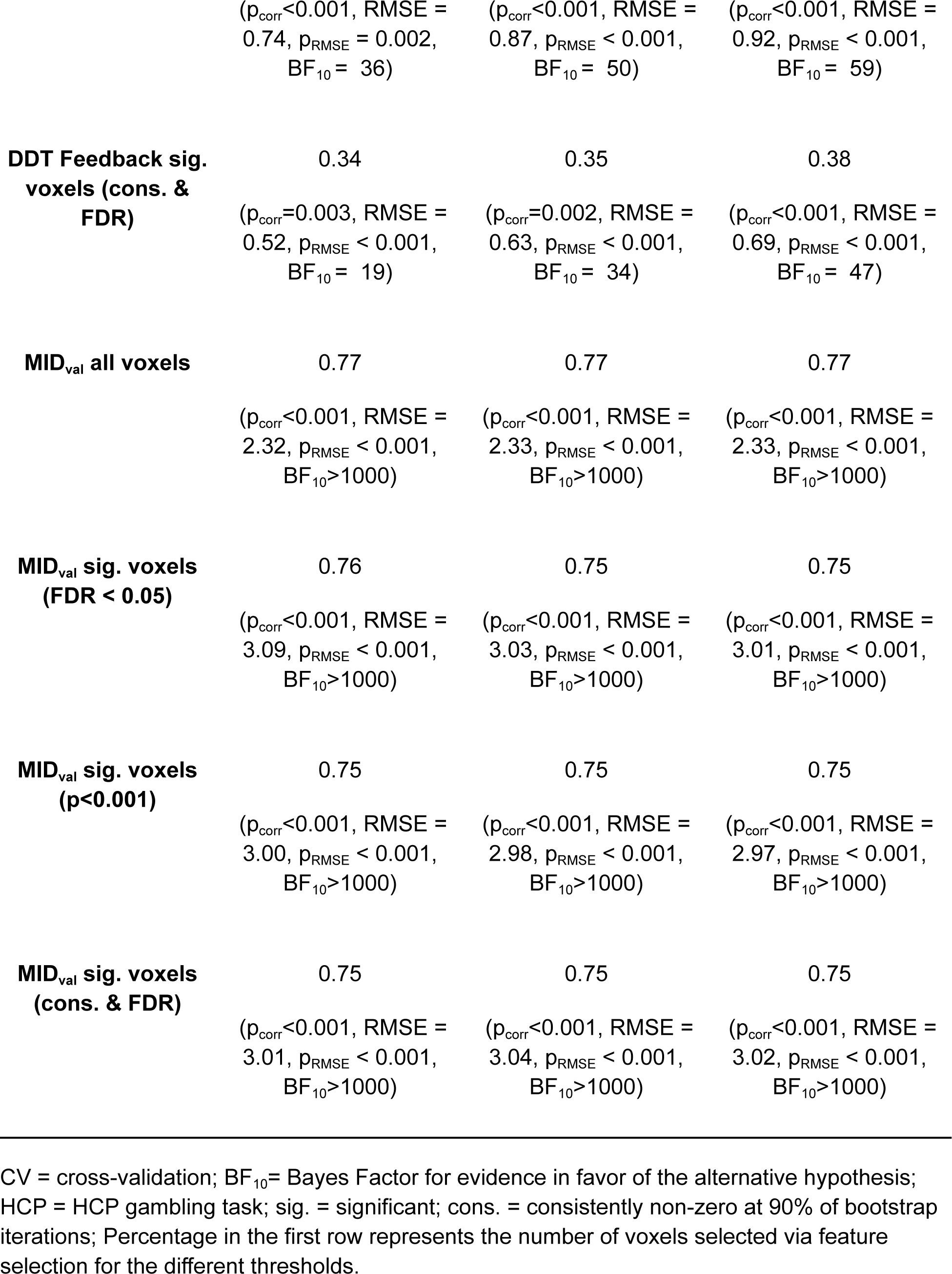
Robustness Check for LASSOPCR on validation tasks.

### Appendix 3: Using Neurosynth masks related to monetary outcomes for feature selection

To compare our data-driven feature selection approach to a more theory driven feature selection approach we also used two Neurosynth maps (Yarkoni et al., 2011; see Table S2) related to monetary outcomes for feature selection within the cross-validation loop. Specifically, we used a meta-analytic map created based on the term *monetary reward* (Association test, FDR corrected for multiple comparisons at p<0.01) and on the term *outcome* (Association test, FDR corrected for multiple comparisons at p<0.01). A similar pattern of results as for the data-driven feature selection approach reported in the main text was found. Again the BRS significantly predicted monetary outcomes in the MID_val_ and the HCP gambling task, but did not significantly predict outcomes in the DDT. Performance on the HCP was slightly higher, whereas performance on the MID_val_ was slightly lower, which was expected as the data-driven feature selection was trained on another version of the MID task which and consequently was more likely to perform higher on a similar task. In contrast, the theory-driven approach was more task independent and more likely to perform similarly well across tasks.

**Figure S1.**
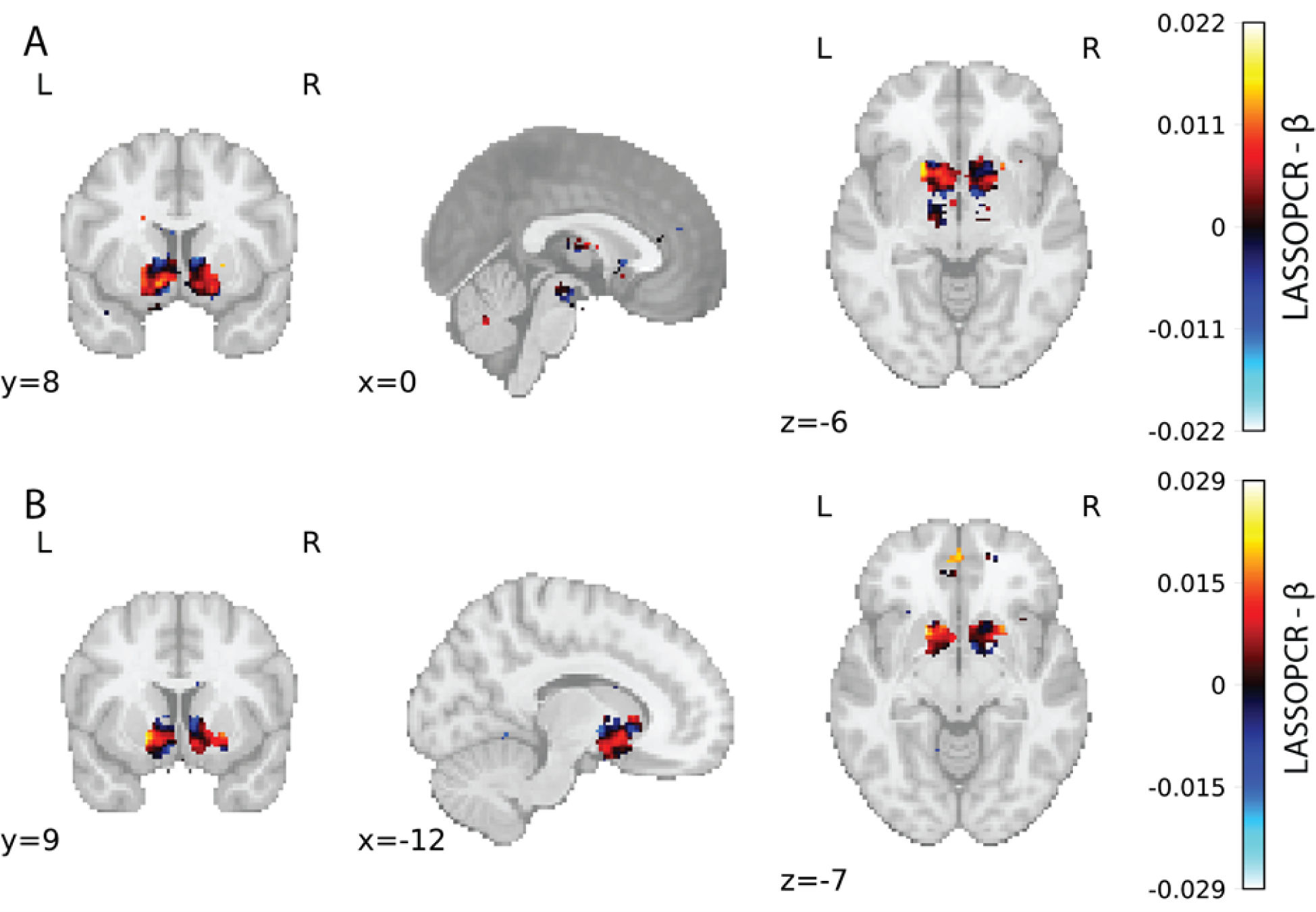
A: Prediction weights derived from the feature-selection approach based on the *monetary reward* meta-analytic map. B: Prediction weights derived from the feature-selection approach based on the *outcome* meta-analytic map.

**Table S2.** Neurosynth maps for monetary reward and outcome

| Network | Studies | Date of | Link to download |
| --- | --- | --- | --- |
| Monetary Reward | 97 | 04.10.2021 | https://neurosynth.org/analyses/terms/monetary%20reward/ |
| Outcome | 385 | 04.10.2021 | https://neurosynth.org/analyses/terms/outcome/ |

**Table S3.**
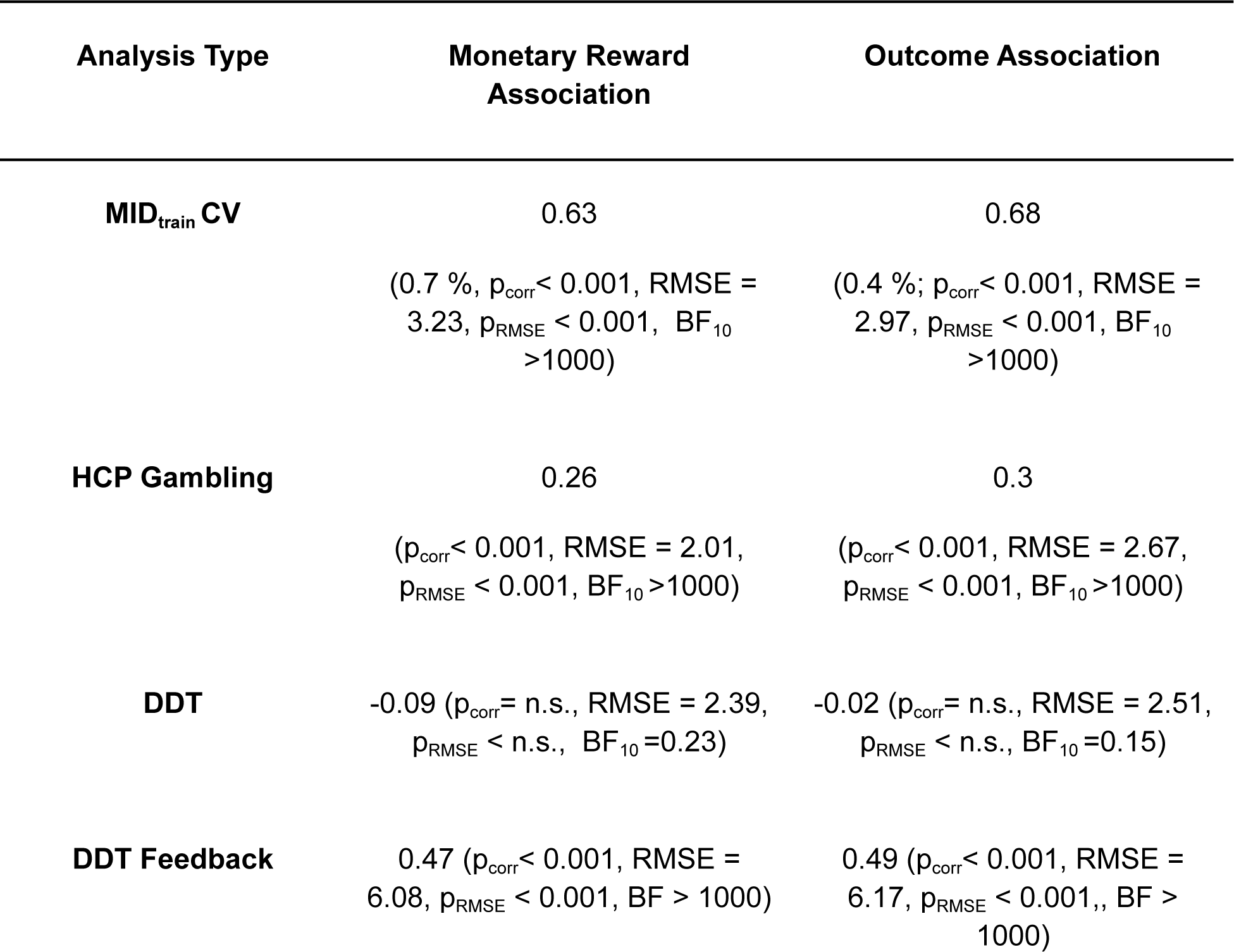
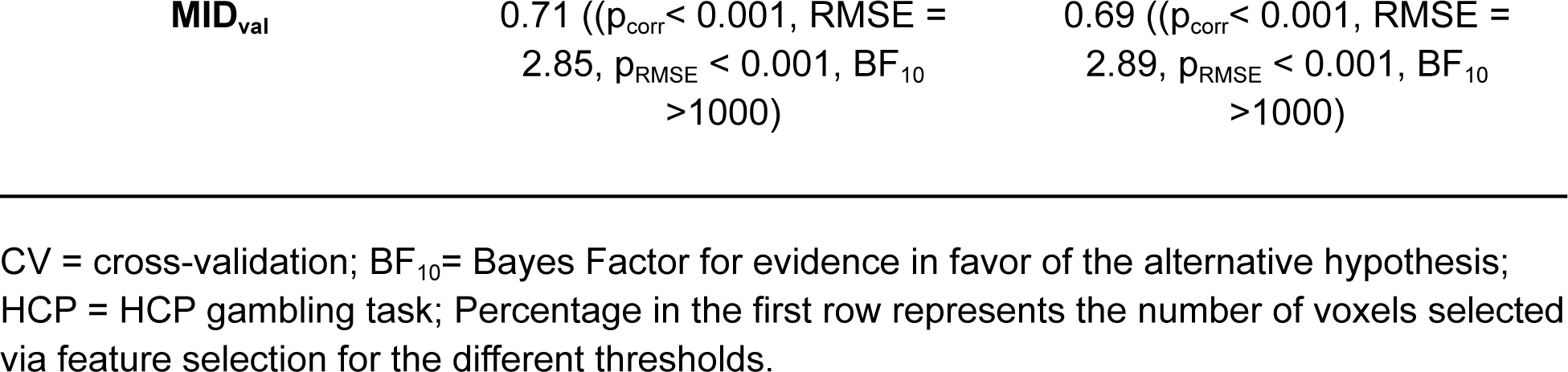
Prediction Performance for Feature Selection based on Neurosynth Maps.

| Analysis Type | Monetary Reward Association | Outcome Association |
| --- | --- | --- |
| **MID<sub>train</sub> CV** | 0.63<br>(0.7 %, $p_{\text{corr}} < 0.001$, RMSE = 3.23, $p_{\text{RMSE}} < 0.001$, $\text{BF}_{10} > 1000$) | 0.68<br>(0.4 %; $p_{\text{corr}} < 0.001$, RMSE = 2.97, $p_{\text{RMSE}} < 0.001$, $\text{BF}_{10} > 1000$) |
| **HCP Gambling** | 0.26<br>($p_{\text{corr}} < 0.001$, RMSE = 2.01, $p_{\text{RMSE}} < 0.001$, $\text{BF}_{10} > 1000$) | 0.3<br>($p_{\text{corr}} < 0.001$, RMSE = 2.67, $p_{\text{RMSE}} < 0.001$, $\text{BF}_{10} > 1000$) |
| **DDT** | -0.09 ($p_{\text{corr}} = \text{n.s.}$, RMSE = 2.39, $p_{\text{RMSE}} < \text{n.s.}$, $\text{BF}_{10} = 0.23$) | -0.02 ($p_{\text{corr}} = \text{n.s.}$, RMSE = 2.51, $p_{\text{RMSE}} < \text{n.s.}$, $\text{BF}_{10} = 0.15$) |
| **DDT Feedback** | 0.47 ($p_{\text{corr}} < 0.001$, RMSE = 6.08, $p_{\text{RMSE}} < 0.001$, $\text{BF} > 1000$) | 0.49 ($p_{\text{corr}} < 0.001$, RMSE = 6.17, $p_{\text{RMSE}} < 0.001$, $\text{BF} > 1000$) |

### Appendix 4: Full List of Clusters for the *BRS* map

**Table S4.** Full list of cluster for the bootstrap thresholded *BRS* map.

| Region | peak_x | peak_y | peak_z | peak_value | volume_mm | nr_voxels |
| --- | --- | --- | --- | --- | --- | --- |
| R Dorsal Striatum | 24 | 14 | -2 | 722.053 | 3456 | 432 |
| L Dorsal Striatum | -20 | 12 | -8 | 97.371 | 3152 | 394 |
| R Occipital Pole | 16 | -92 | -8 | 679.417 | 1464 | 183 |
| vmPFC | 2 | 44 | -4 | 576.394 | 1248 | 156 |
| L Frontal Pole | -30 | 62 | 0 | 454.149 | 152 | 19 |
| L Precuneus | -2 | -52 | 52 | -467.939 | 128 | 16 |
| R Occipital Pole | 20 | -102 | -4 | 425.175 | 112 | 14 |
| R Supramarginal Gyrus | 62 | -36 | 42 | -442.408 | 96 | 12 |
| Left Fusiform Cortex | -46 | -62 | -22 | -449.661 | 96 | 12 |
| R Occipital Pole | 16 | -90 | 4 | -454.105 | 96 | 12 |
| L Dorsal Striatum | -30 | -12 | 6 | 466.043 | 88 | 11 |
| R Precentral Gyrus | 32 | -22 | 56 | 407.251 | 80 | 10 |
| R SMA | 14 | 14 | 64 | -465.555 | 72 | 9 |
| Left Occipital Pole | -24 | -92 | -8 | 388.064 | 72 | 9 |
| L SMA | -8 | 24 | 64 | -463.509 | 64 | 8 |
| L Postcentral Gyrus | -48 | -20 | 42 | -418.744 | 64 | 8 |
| R Postcentral Gyrus | 44 | -28 | 54 | 458.414 | 56 | 7 |
| R MTG | 62 | -38 | -2 | -401.238 | 56 | 7 |
| R MTG | 56 | -30 | -4 | -392.721 | 48 | 6 |
| L Frontal Pole | -22 | 58 | -4 | 420.486 | 48 | 6 |
| L Postcentral Gyrus | -62 | -8 | 18 | -371.324 | 48 | 6 |
| L Frontal Pole | -10 | 64 | 4 | 464.849 | 48 | 6 |
| L Frontal Pole | -22 | 64 | 4 | 409.869 | 48 | 6 |
| L MTG | -54 | -26 | -8 | -452.319 | 48 | 6 |
| Left Occipital Pole | -6 | -92 | 0 | -47.522 | 40 | 5 |
| L ACC | -6 | 40 | -2 | 374.225 | 40 | 5 |
| L Cerebellum | -22 | -58 | -20 | 458.105 | 40 | 5 |
| R ACC | 4 | 10 | 32 | 358.943 | 40 | 5 |
| L Frontal Pole | -20 | 62 | 2 | 425.523 | 40 | 5 |
| L Postcentral Gyrus | -32 | -30 | 64 | -407.198 | 32 | 4 |
| Precuneus | -6 | -62 | 54 | -366.421 | 32 | 4 |
| Precuneus_L | -10 | -64 | 66 | -362.473 | 24 | 3 |
| Left Occipital Pole | -14 | -96 | -10 | 407.336 | 24 | 3 |
| Supp_Motor_Area_R | 2 | 18 | 60 | -396.662 | 24 | 3 |
| Frontal_Sup_2_L | -28 | 52 | -4 | 401.706 | 24 | 3 |
| R SMA | -36 | -32 | 66 | -403.352 | 24 | 3 |
| L Caudate Nucleus | -8 | 18 | -2 | 366.792 | 24 | 3 |
| vmPFC | -6 | 48 | -8 | 413.769 | 24 | 3 |
| L Intracalcarine Cortex | -8 | -88 | 4 | -417.215 | 24 | 3 |
| L Lateral Occipital Cortex | -22 | -70 | 40 | -41.666 | 24 | 3 |
| R Supramarginal Gyrus | 68 | -28 | 34 | -372.851 | 16 | 2 |
| R IFG | 58 | 30 | 20 | -370.708 | 16 | 2 |
| R Temporal Pole | 52 | 10 | -24 | -401.481 | 16 | 2 |
| L Dorsal Striatum | -28 | -8 | 12 | 366.682 | 16 | 2 |
| R MTG | 66 | -46 | 6 | -3.464 | 16 | 2 |
| R Lateral Occipital Cortex | 34 | -86 | 28 | -413.061 | 16 | 2 |
| R Lateral Occipital Cortex | 32 | -78 | 30 | -369.657 | 16 | 2 |
| R Superior Temporal Gyrus | 64 | 0 | -6 | 396.061 | 16 | 2 |
| L Occipital Pole | -16 | -98 | -10 | 395.429 | 16 | 2 |
| R Occipital Pole | 50 | -26 | 58 | 347.575 | 16 | 2 |
| L MTG | -44 | -62 | 20 | -342.001 | 16 | 2 |
| L Parietal Operculum Cortex | -36 | -26 | 18 | -48.481 | 16 | 2 |
| R Lateral Occipital Cortex | 18 | -74 | 52 | -337.939 | 16 | 2 |
| L Lingual Gyrus | -8 | -86 | -14 | -391.361 | 16 | 2 |
| L Frontal Pole | -6 | 66 | 8 | 364.597 | 16 | 2 |
| R Frontal Pole | 16 | 62 | 28 | -358.984 | 16 | 2 |
| R Superior Frontal Gyrus | 12 | 32 | 62 | -375.761 | 16 | 2 |
| R Intracalcarine Cortex | 8 | -88 | 2 | -361.447 | 16 | 2 |

L = Left ; R = Right; vmPFC = ventromedial prefrontal cortex. SMA = supplementary Motor Area; MTG = Middle Temporal Gyrus; ACC = Anterior Cingulate Cortex; IFG = Inferior Frontal gyrus; Only clusters of at least 2 voxels are shown, a complete list can be found in Supplementary Table xx. All voxels were used in the analysis. The table was generated using the python package Atlasreader (Notter et al., 2019).

